# Epigenetic Profiling of Human Insulinomas Reveals AP-1 Family as Critical Regulators of Beta Cell Maturation

**DOI:** 10.64898/2026.05.29.728759

**Authors:** Xuedi Wang, Saul Carcamo, Luca Lambertini, Peng Wang, Hongtao Liu, John D. Paulsen, Rachel Brody, Gustavo Fernandez-Ranvier, Dan Hasson, Andrew F. Stewart, Esra Karakose

**Author notes:** indicates lead and corresponding contact Esra Karakose PhD, Assistant Professor of Medicine, Diabetes Obesity and Metabolism Institute The Icahn School of Medicine at Mount Sinai Atran 5, PO Box 1152, One Gustav Levy Place New York, NY, 10029.

## Abstract

Insulinomas are rare and benign human pancreatic adenomas that overproduce insulin and display increased beta cell mass. We and others have shown that transcriptomic and genomic profiling on insulinomas provides a data mine for identifying targets that can be manipulated to induce human beta cell regeneration. Majority of causative genetic variants in insulinomas involve epigenetic regulatory genes. Yet, specifically how these variants lead to human beta cell expansion and increased function is largely unknown. Here, we performed bulk and single-nucleus epigenomic and transcriptomic profiling to define regulatory alterations in human insulinomas. Bulk ATAC-seq and H3K27Ac ChIP-seq revealed significant enrichment of AP-1 transcription factor binding motifs within beta cell-associated open chromatin/enhancer regions in normal islets, accompanied by robust expression of AP-1 family members; in contrast, insulinomas exhibited marked reductions in both AP-1 motif enrichment and AP-1 expression. Our snRNA-seq and snATAC-seq profiling across four independent insulinomas identified a consistent and previously unrecognized signature defined by suppression of AP-1 transcription factors and widespread loss of chromatin accessibility at AP-1 binding sites, particularly at enhancers governing beta cell identity. Collectively, these results establish AP-1-mediated regulatory programs as critical determinants of beta cell maturation and define their disruption as a signature among human insulinomas.

## Introduction

Insulinomas are rare pancreatic neoplasms. Their incidence is estimated to be 1-3 cases per million people per year^1–3^. Insulinomas classically produce dramatic and dangerous hypoglycemia that results from inappropriate oversecretion of insulin, causing recurrent episodes of sudden reduced consciousness, often accompanied by accidents and injury^1–3^. This leads to assessment of blood glucose and insulin, and diagnostic imaging studies that identify the causative pancreatic tumor, which is then removed surgically.

Insulinomas are a subgroup of a larger family of neuroendocrine neoplasms (NENs) that may occur in multiple organs, including the pancreas, stomach, intestine, lung and elsewhere^1–3^. When they occur in the pancreas, they are referred to as pancreatic neuroendocrine tumors (PNETs). PNETs may be benign or malignant, and may secrete multiple different peptide hormones [e.g., insulin, glucagon, adrenal corticotrophic hormone (ACTH), corticotrophin-releasing hormone (CRH), parathyroid hormone-related protein (PTHRP), and calcitonin, among others] as well as bioactive small molecules such as serotonin^1–3^. Depending on the hormone produced, NENs and PNETs may be associated with specific clinical syndromes, exemplified by hypoglycemia (insulinomas), diabetes (glucagonomas), hypercortisolism (ACTH- and CRH-producing NENs), hypercalcemia (PTHrP-producing NENs), etc. Some NENs produce no identifiable hormones and have no associated paraneoplastic syndrome, and are considered “non-functional” NENs. Conversely, a single NEN can simultaneously produce multiple different hormones and clinical syndromes^1–3^.

Insulinomas, like other NENs and PNETs may be classified pathologically as benign or malignant based on pathological features or histologic aggressiveness, local or distant tissue invasion, and tumor cell proliferation rate, assessed by the immunohistochemical labeling of tumor cells with the proliferation marker Ki-67^1–3^. Most are classified as benign, based on Ki-67 labeling indices below 3%^1^. In terms of molecular etiology, mutations in multiple genes have been reported in human insulinomas and other PNETs, the two most common being inactivating mutations in *MEN1*, encoding the menin protein, a member of the Trithorax or COMPASS family of epigenetic regulators, and the YY1 transcription factor, encoded by *YY1*, another epigenetic regulator^1–9^. Strikingly, the majority of insulinomas that do not have defects in *MEN1* or *YY1*, display structural and/or copy number variants (CNVs) in other epigenetic regulatory proteins, exemplified by KDM6A (UTX), KMT2A (MLL1), KNT2D (MLL2), KMT2C (MLL3), JARID2, EZH2, ARID5B, BMI1, EED, CREBBP(CBP) and others^5,7,8,10^. Bulk RNA-sequencing and proteomics on human insulinomas confirms and extends these observations^7,8,10^. In contrast to benign insulinomas, the genomic and epigenetic landscape in malignant PNETs and insulinomas includes the above, but also another set of commonly recurring variants in the cell death and de-differentiation factor, DAXX, and the epigenetic transcriptional regulator, ATRX^10–12^. Finally, it is usual to observe multiple mutations in multiple of the epigenetic regulatory genes described above.

Here, we describe detailed bulk ATAC-seq and H3K27Ac ChIP-seq on human insulinomas and compare these to normal pancreatic islets. Unexpectedly, we observe, significant motif enrichment for AP-1 (Jun and Fos) family transcription factor binding sites in beta cell-associated open chromatin/enhancer regions in normal pancreatic beta cells. In contrast, we observe that these AP-1 binding sites and expression of AP-1 members are significantly reduced in insulinomas. The AP-1 factors in question include the Jun and Fos families, as detailed below.

Because of their rarity, variable presentations and multiple subtypes, as of this writing, no studies have assessed the epigenetic landscape of insulinomas at the single cell level. Therefore, we also describe single nucleus RNA-seq and ATAC-seq in four separate human insulinomas and compare these to normal pancreatic islets. As anticipated, we identify an altered transcriptomic and epigenetic landscape within the beta cells of insulinomas as compared to normal beta cells. Most importantly, the single cell approaches confirm and extend the presence of a strong and previously unrecognized signature involving: 1) suppression of expression of the AP-1 family of transcription factors; and, 2) widespread loss of chromatin accessibility for AP-1 factors, particularly enriched at beta cell-relevant enhancers. Collectively, these findings demonstrate that AP-1 factors play a crucial role in defining beta cell maturation and functional differentiation, and their loss and/or disruption of binding opportunities chromatin-wide provide a characteristic signature of human insulinomas.

## Results

### Bulk Epigenetic and Transcriptome Profiling of Insulinomas Nominates AP-1 Family Transcription Factors as Critical Regulators of Beta Cell Phenotype

We performed bulk ATAC-seq and H3K27Ac ChIP-seq to comprehensively characterize the active cis-regulatory landscape of beta cells and insulinomas. These analyses were conducted on four insulinoma samples and compared to three FACS-purified pancreatic beta cell preparations (**Suppl. Fig.1A-B**). Differential accessibility analysis and differential H3K27ac enrichment analysis was performed on ATAC-seq and H3K27Ac ChIP-Seq data respectively (**Suppl. Table 1,2**). To systematically define regulatory elements, we first identified ATAC-seq peaks that show increased accessibility in beta cells, increased accessibility in insulinomas, or static, and then evaluated their H3K27Ac status (differential enrichment, static, or absence of H3K27Ac signal). This two-feature integration approach initially yielded eight distinct peaks’ clusters **(Suppl. Fig. 1C)**, which were merged into three major clusters based on the concordance of chromatin accessibility and H3K27Ac signals across samples (**Fig. 1A**). Cluster 1 (8,455 peaks) represents beta-cell-associated cis-regulatory regions, defined by preferential chromatin accessibility and/or H3K27Ac enrichment in beta cells. Cluster 2 (1,620 peaks) represents insulinoma-associated cis-regulatory regions, defined by concordant increases in both chromatin accessibility and H3K27Ac signal in insulinoma samples. Cluster 3 (21,102 peaks) represents shared cis-regulatory regions, comprising of genomic regions without differential activity in either.

**Figure 1:**
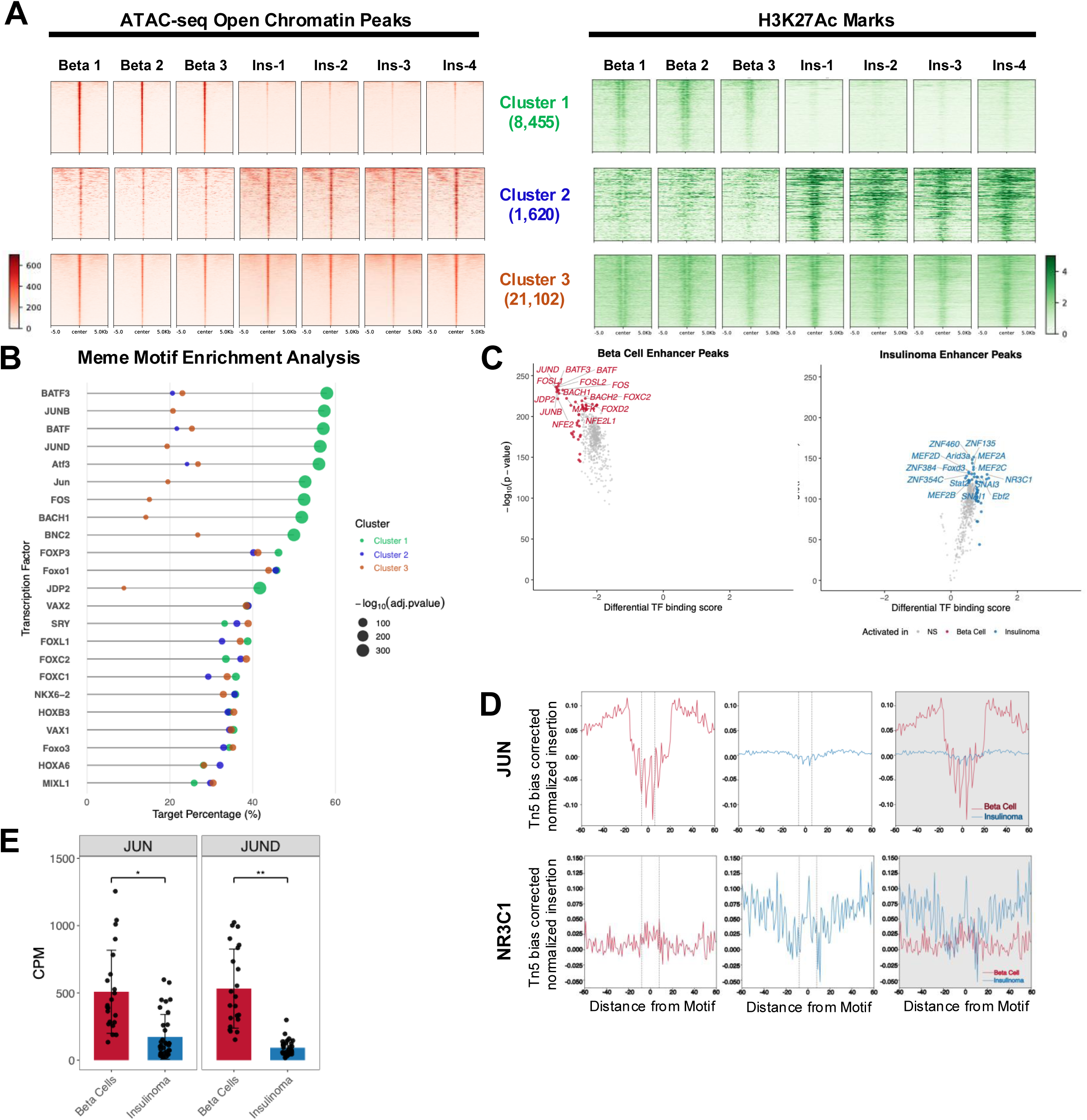
Bulk Epigenetic and Transcriptome Profiling of Insulinomas Nominates AP-1 Family Transcription Factors as Critical Regulators of Beta Cell Phenotype. **A.** Heatmaps of normalized ATAC-seq peaks (left) and corresponding H3K27ac signal (right) obtained from 3 normal beta cell donors and 4 insulinoma patients. Heatmap is centered around ATAC peaks (Cluster 1 beta cells n = 8,455 ATAC peaks; Cluster 2 Insulinomas n = 1,620; and cluster 3 shared n = 21,102 peaks) and ranked by ATAC signal intensity. **B**. Lollipop plot showing the top 10 significantly enriched transcription factor binding motifs for each cluster. Dot size indicates statistical significance; the x-axis shows the percentage of input peaks containing each motif. **C**. Volcano plots showing TOBIAS differential transcription factor footprinting results for cluster 1 (left) and cluster 2 (right). Transcription factors in the top 5% by significance and magnitude of binding score change are highlighted. The top 15 most significant transcription factors are labelled for each group. The x-axis shows the differential binding score for indicated peaks. **D.** Transcription factor occupancy footprinting for JUN (n = 1,773 motif sites) (top), and NR3C1 (n = 248 motif sites) (bottom). Plots show aggregated, Tn5 insertion-bias-corrected ATAC-seq signal (y-axis) centered on a 60 bp window around the motif center. **E**. Bar chart of bulk RNA-seq expression (CPM) for AP-1 genes *JUN* and *JUND* from differential gene expression analysis of 22 FACS-sorted beta cells (red) versus 34 insulinomas (blue). Bars indicate the group mean (±SD); dots represent individual samples. Brackets denote the between-group comparison; significance is shown as *FDR < 0.05 and **FDR < 0.01.

KEGG Pathway enrichment analysis for each regulatory cluster (**Suppl. Fig. 2A, Suppl. Table 3**) revealed distinct biological processes associated with each regulatory cluster. Notably, the beta cell-associated cluster (Cluster 1) was significantly enriched for the TGF-β signaling pathway. This observation is consistent with prior studies from ourselves and others demonstrating that inhibition of TGF-β signaling promotes proliferation in otherwise quiescent human pancreatic beta cells^13,14^. In contrast, the shared cluster (Cluster 3) showed strong enrichment for pathways related to maturity-onset diabetes of the young (MODY), reflecting preservation of beta cell differentiation programs in both normal and insulinoma contexts.

We next performed transcription factor (TF) motif enrichment using MEME. We found transcription factor motifs such as AP-1 to be the dominant regulator of chromatin accessibility in beta cell-specific enhancers (Cluster 1), whereas FOX and HOX family motifs were enriched in the insulinoma-enhancers although to a lesser extent (Cluster 2) (**Fig. 1B, Suppl. Table 4**). To further determine transcription factor motif regulators, we performed TF footprinting analysis, and found that insulinomas have markedly reduced binding at AP-1 sites when compared to beta cells. (**Fig. 1C-D; Suppl. Fig. 2B, Suppl. Table 5**). In contrast, insulinoma-associated regions (Cluster 2) showed binding of the glucocorticoid receptor NR3C1 and the ZNF family TFs (**Fig. 1C-D; Suppl. Fig. 2C, Suppl. Table 5**).

We reasoned that the accessibility of a given transcription factor motif should correlate with the expression of the associated transcription factor in beta cells and insulinomas. To further predict causative regulators of motif accessibility, we integrated our DNA binding motif and footprinting analyses with our previously published bulk RNA-seq expression data on 37 insulinomas vs. 22 sets of FACS-sorted beta cells^7^. Using this approach, we found a strong correlation of motif usage and expression of select AP-1 members, specifically *JUN* and *JUND* in beta cells compared to insulinomas (**Fig. 1E),** and their binding partners FOS and FOSB **(Suppl. Fig. 2B).** Notably, *JUN* and *JUND* were the only AP-1 family members that met our criteria for both high expression (CPM > 90) and significant differential downregulation in insulinomas (**Fig. 1E**). Other AP-1 family TFs exhibited either low expression levels or were not differentially expressed between conditions (**Suppl. Fig. 3C**). Collectively, these data nominate the AP-1 family of transcription factors as key regulators of the beta cell phenotype.

### Single Cell Approaches Identify Distinct Subtypes of Beta Cells in Insulinomas

Analysis to this point were performed on bulk insulinoma RNAseq datasets. Since insulinomas contain many cell types (vascular cells, immune cells, mesenchymal cells, pancreatic acinar cells, etc.), to further achieve cell type-resolved characterization of these regulatory programs and to understand whether the changes we observed occur specifically in insulinoma beta cells, we next performed single-nucleus RNA-seq (snRNA-seq) and snATAC-seq on insulinomas. We performed snRNA-seq on three insulinoma samples, and identified multiple cell types, including endocrine, acinar, endothelial, immune, and delta cells, as well as four transcriptionally distinct beta cell types (**Suppl. Fig. 4A,B, Suppl. Table 6**). Notably, one of these beta cell populations was characterized by a proliferative signature, representing a minor but distinct population of proliferating beta cells (**Suppl. Fig. 4A,B, Suppl. Table 7**). Consistent with observations from bulk transcriptomic and genomic sequencing analyses^5,7,8,10,11^, and pathology analysis of insulinomas^1–3^, we detected substantial inter-sample heterogeneity among the insulinomas. For instance, sample 3 contributed to more than 50% of the proliferative beta cell subcluster and over 25% were found in sample 2, whereas sample 1 only contributed approximately 1% of total proliferative beta cell population (**Suppl. Fig. 5A,B**). Among all annotated cell types in the snRNA-seq data, only the endocrine compartment exhibited a comparable cell abundance distribution across all samples (**Suppl. Fig. 5B**), as expected because the surgically excised insulinoma sample was the principal tissue submitted for snRNA-seq.

We next conducted single-nucleus ATAC sequencing (snATAC-seq) on four insulinoma samples, three of which overlapped with the snRNA-seq sample set. Chromatin accessibility profiling revealed six distinct beta cell subclusters (Beta_1–Beta_6) (**Fig. 2A–C, Suppl. Table 8**), in addition to acinar, ductal, endothelial, microglial, and stellate cell populations (**Fig. 2A–C, Suppl. Table 9).** The identification of cluster-specific accessible chromatin regions specific to individual beta cell subclusters (**Fig. 2B,C**) further underscores the accessible chromatin landscape heterogeneity in insulinoma beta cells. Similarly to the transcriptomic data, the snATAC-seq dataset demonstrated pronounced heterogeneity, with certain beta cell subclusters (e.g., Beta_2 and Beta_3) predominantly or exclusively deriving from individual samples (**Suppl. Fig. 5C,D).** Genomic annotation of open chromatin regions across all cell types showed that the majority of peaks (over 85%) were located in distal regulatory elements (**Fig. 2D**), consistent with the enhancer-centric regulatory landscape found in our bulk ATAC-seq and H3K27Ac ChIP-seq data (**Fig. 1A**).

**Figure 2:**
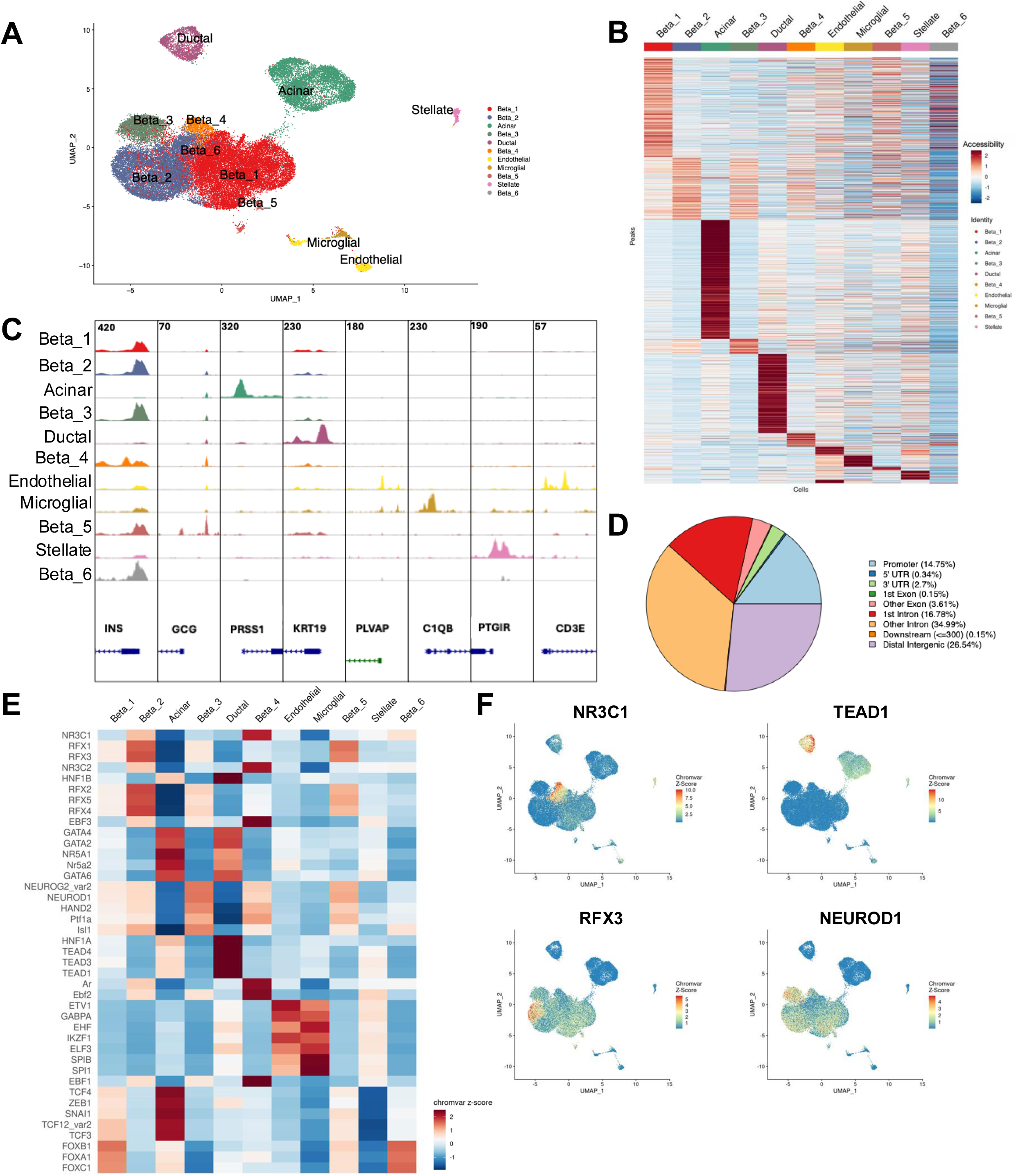
Single Cell Approaches Identify Distinct Subtypes of Beta Cells in Insulinomas. **A.** UMAP of integrated insulinoma snATAC-seq data (n=4; 45,923 cells in total) colored by inferred cell type clusters. **B.** Heatmap showing z-scores of normalized accessibility of 20,000 sub-sampled cluster specific peaks. **C.** Capture of the genome browser tracks showing aggregated chromatin accessibility profiles for each cell cluster at canonical marker gene loci used for cell type annotation. Numbers on top of each column indicate the y-axis value for the given locus. **D.** Genomic distribution of 214,334 peaks called from the integrated snATAC-seq object. Promoters are defined as ±1,000 bp from the TSS. **E.** Heatmap of scaled chromVAR motif deviation scores for the top 5 discriminating transcription factor motifs identified from each cell cluster versus all other clusters. **F.** UMAPs as in A colored by chromVAR deviation scores for NR3C1, TEAD1, RFX3, and NEUROD1. Color scale boundaries were set at the 1st and 99th percentiles of the deviation score distribution.

Finally, transcription factor (TF) motif activity analysis revealed cell type- and subtype-specific regulatory programs within the snATAC-seq dataset (**Fig. 2E, Suppl. Table 11**). For example, NR3C1 activity was enriched in the Beta_4 subcluster, TEAD1 activity was prominent in ductal cells, and RFX3 activity was elevated in the Beta_2 subcluster. In contrast, NEUROD1 exhibited consistently high activity across all beta cell subtypes (**Fig. 2F**).

### Comparative Single-Cell Analysis of Normal Islets and Insulinomas Reveals Altered AP-1 Transcriptional Programs in Beta Cells

To systematically compare our insulinoma single-nucleus datasets with independently reported normal human islets, we used a previously published multiome dataset, from Wang et al^15^, comprising six healthy human islet donors (GEO: GSE200044). These data were processed and integrated using an identical analytical pipeline to that employed to our insulinoma datasets to ensure methodological consistency. Analysis of this reference dataset identified approximately 20,000 cells, of which ∼50% were annotated as beta/endocrine cells (**Suppl. Fig. 6A,B).** Our integrated insulinoma snRNA-seq dataset comprised approximately 10,000 total cells, including ∼8,000 annotated beta/endocrine cells (**Suppl. Fig. 6C**). The corresponding snATAC-seq dataset included ∼46,000 nuclei, of which ∼35,000 were classified as beta cells (**Suppl. Fig. 6D**). The enrichment of beta/endocrine cells in insulinomas is expected since the majority of the samples were surgically excised insulinoma tissue.

To enable direct cell type-resolved comparisons, we next projected insulinoma snRNA-seq profiles onto the normal islet reference dataset. This mapping approach demonstrated that the majority of insulinoma cells aligned with the beta cell identity in the reference dataset (**Fig. 3A**). A similar projection of insulinoma snATAC-seq profiles yielded concordant results, with most cells also mapping to beta cell annotations (**Fig. 3B**). These projection-derived labels were subsequently used for downstream comparative analyses between insulinoma and normal islet beta cells.

**Figure 3:**
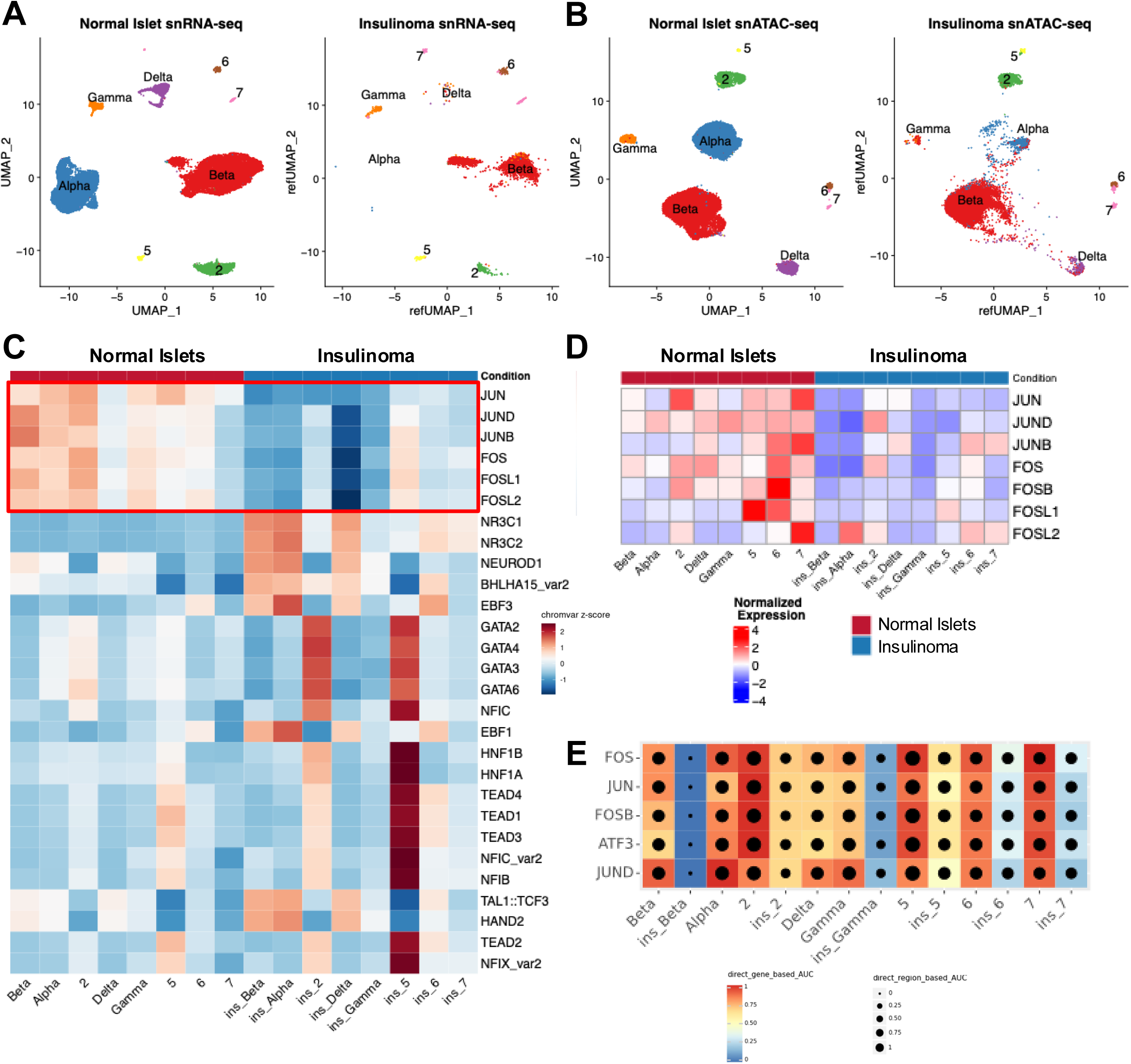
Comparative Single-Cell Analysis of Normal Islets and Insulinomas Reveals Altered AP-1 Transcriptional Programs in Beta Cells. **A.** UMAP of integrated normal islet snRNA-seq data (n = 6; GSE200044) colored by inferred cell type clusters (left), and UMAP showing the MapQuery projection of insulinoma snRNA-seq data (n = 3) onto the normal islet reference (right). **B.** UMAP of integrated normal islet snATAC-seq data (n = 6; GSE200044) inferred cell type clusters (left), and UMAP showing the MapQuery projection of insulinoma snATAC-seq data (n = 4) onto the normal islet reference (right). **C.** Heatmap of scaled chromVAR motif deviation scores for the top 5 enriched transcription factor motifs per cell type, comparing insulinoma to matched normal islet cell types. Cell type clusters are defined by projection of insulinoma cells onto the normal islet reference as shown in (B). **D.** Heatmap of z-score of normalized gene expression for AP-1 family members. Data is scaled across clusters. Cell type clusters are defined by projection of insulinoma cells onto the normal islet reference as shown in (A). **E.** Heatmap/dot-plot showing the scaled AUC scores for canonical AP-1 activator eRegulons from Suppl Fig. 7. Heatmap color indicates the expression-based AUC of target genes; dot size indicates the AUC of associated regulatory regions.

To interrogate transcription factor (TF) regulatory landscapes in the insulinoma samples, we performed ChromVAR analysis between matched cell types from both conditions (**Fig. 3C, Suppl. Table 11**). Comparison of normal islet beta cells versus beta cells from insulinomas identified increased TF activity for the AP-1 family TF family (including *JUN*, *JUND*, *JUNB*, *FOS*, and *FOSL1*) in beta cells from normal islets, whereas motif accessibility was markedly reduced in insulinoma-derived beta cells (**Fig. 3C**). Consistent with these findings, snRNA-seq data demonstrated higher expression of *JUN*, *JUND*, and *FOS* in normal islet beta cells relative to insulinoma beta cells (**Fig. 3D).** Conversely, NR3C1 TF activity was preferentially enriched in insulinoma beta, alpha, and delta cells compared to their normal islet counterparts, consistent with our bulk ATAC-seq findings. In addition, TEAD family motifs showed preferential activity in exocrine cell populations in insulinomas, suggesting dysregulation of Hippo-YAP/TAZ signaling in the exocrine compartment (**Fig. 3C**).

To move beyond motif activity and to further reconstruct the gene regulatory networks in normal islets and insulinomas, we performed SCENIC+ analysis on the integrated insulinoma and normal islet datasets. To enable joint analyses of both snRNA-seq and snATAC-seq, pseudo-multiome meta cells were generated by SCENIC+, and cell types with insufficient representation in either modality, including alpha cells and delta cells in insulinomas, were excluded from this step and thus are absent in the regulon analysis. Similar to our ChromVAR analysis, we found increased regulon activity of ZNF385D, a beta-cell transcription factor marker gene^16^ and AP-1 family regulons including JUN, JUND, FOS, and FOSB, in beta cells from normal islets compared to insulinomas (**Fig. 3C, Suppl. Fig. 7**). Transcription factors that maintain beta cell identity, including MAFA and PDX1, showed high regulon activity in both normal islet and insulinoma beta cells, suggesting that core beta cell identity is preserved in insulinomas. In contrast, insulinoma beta cells showed preferential activity of RB1, KLF12, and NCALD regulons (**Suppl. Fig. 7**). RB1 has been shown to play an important role in pancreatic beta cell biology by stabilizing PDX1 and promoting beta cell differentiation^17^ and maintaining beta cell quiescence^18–20^.

Collectively, these results provide convergent evidence across chromatin accessibility, gene expression, and regulatory network inference analyses that AP-1 transcriptional programs are significantly diminished in insulinoma beta cells relative to beta cells from normal human islets.

### AP-1 Family Transcription Factors Regulate Beta Cell Proliferation and Differentiation

To model the downregulation of AP-1 TFs observed in insulinoma beta cells and to assess their functional role in human beta cell biology, we performed targeted silencing of *JUN* and *JUND* in primary human cadaveric islets (**Suppl. Fig. 8A,C**). Silencing of *JUND* resulted in a modest compensatory upregulation of *JUN* and *JUNB* (**Suppl. Fig. 8B**). In contrast, *JUN* silencing led to a significant decrease in *JUND* and *JUNB* expression (**Suppl. Fig. 8D**), suggesting that *JUN* may act as a central regulator within the AP-1 transcriptional network.

We next examined the effect of *JUN* and *JUND* silencing on expression of key cell cycle regulators, including cyclins, cyclin-dependent kinases (CDKs), and CDK inhibitors (CDKIs). Notably, *JUN* silencing resulted in broad downregulation of CDKIs, whereas *JUND* silencing had comparatively modest effects on cell cycle gene expression (**Fig. 4A,B**). Consistent with prior findings demonstrating that significantly reduced *CDKN1C-p57* expression promotes beta cell proliferation, *JUN* silencing led to a small but reproducible increase in the proportion of Ki67-positive beta cells, indicating a modest enhancement of proliferative activity (data not shown). In contrast, *JUND* silencing had minimal impact on beta cell proliferation. Analysis of cyclin and CDK expression further revealed that *JUN* silencing significantly downregulated the expression of *CCNA1*, *CCNB1*, *CCNB2*, *CCND3* and *CDK4* and increased expression of *CCNA2* and *CCNE1*, whereas *JUND* silencing produced only minor changes (**Fig. 4C,D**). These observations suggest that *JUN* may serve as the major regulator of cell cycle activity among the AP-1 family members in human beta cells.

**Figure 4:**
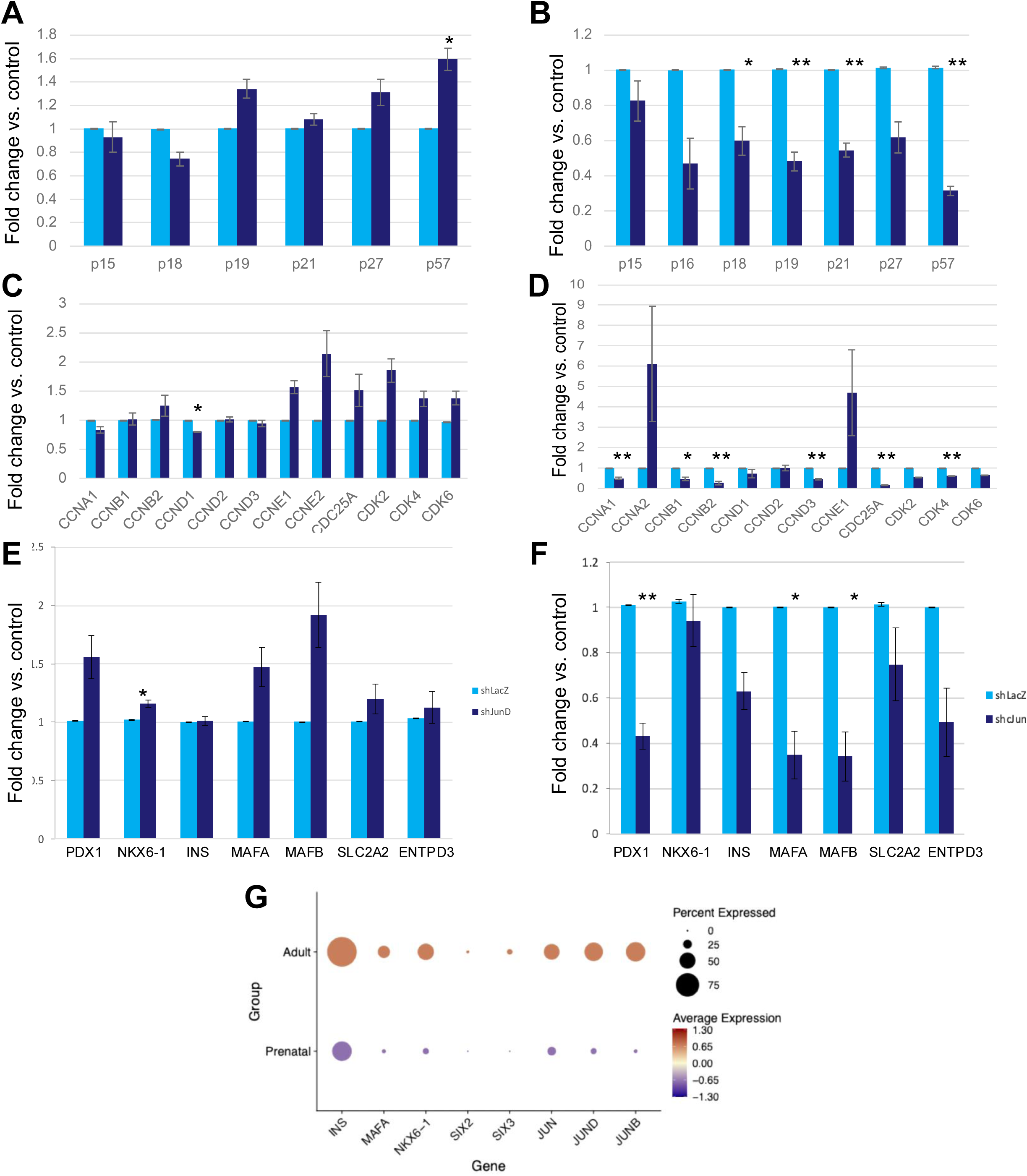
AP-1 Family Transcription Factors Regulate Beta Cell Proliferation and Differentiation. qRT-PCR results from dispersed human islets infected with small hairpin RNA targeting *JUN* or *JUND*. **A.** Gene expression levels for Cyclin-dependent Kinase Inhibitors upon *JUND* silencing. **B.** Gene expression levels for Cyclin-dependent Kinase Inhibitors upon *JUN* silencing. **C.** Gene expression levels for Cyclins and CDKs upon *JUND* silencing. **D.** Gene expression level for Cyclins and CDKs upon *JUN* silencing. **E.** Gene expression levels for beta cell differentiation marker genes upon *JUND* silencing. **F.** Gene expression levels for beta cell differentiation marker genes upon *JUN* silencing. **G.** Dot Plot showing average expression for key endocrine/beta-cell genes (*INS*, *MAFA*, *NKX6-1*, *SIX2*, *SIX3*) and AP-1 genes (*JUN*, *JUND*, *JUNB*), in integrated scRNA-seq from publicly available datasets. Dot size indicates the percentage of cells expressing each gene, and dot color indicates average scaled expression within each group.

We next assessed the impact of AP-1 TF perturbation on beta cell identity by analyzing the expression of canonical beta cell marker genes. Silencing of *JUN* resulted in a significant downregulation of key beta cell markers, including *PDX1*, *MAFA* and *MAFB*, whereas *JUND* silencing led to a significant upregulation in *NKX6-1* and a modest upregulation of *PDX1*, *MAFA*, and *MAFB* (**Fig. 4E,F**). Collectively, these findings suggest that AP-1 family members regulate beta cell proliferation and differentiation, with *JUN* playing a particularly critical role in maintaining beta cell identity.

To further contextualize these findings in a developmental context, we analyzed single-cell datasets comprising human islets from adult and prenatal donors. Expression of *JUN*, *JUND*, and *JUNB* was markedly reduced in prenatal beta cells, mirroring the expression patterns of established beta cell maturity markers such as *INS*, *MAFA*, *NKX6-1*, *SIX2*, and *SIX3* (**Fig. 4G; Suppl. Fig. 9A-C**). Together, these data support a role for AP-1 TFs in the regulation of beta cell maturation and differentiation.

### AP-1 Transcription Factors Regulate Beta Cell Gene Expression via Enhancer Elements

To elucidate the mechanisms by which AP-1 TFs regulate beta cell gene expression, we first examined AP-1 motif enrichment at the promoters of beta cell marker genes. However, AP-1 motifs were largely absent from promoter regions, leading us to investigate whether AP-1 TFs perform their regulatory role through distal regulatory elements. We therefore performed peak-to-gene linkage analysis to identify distal regulatory elements whose chromatin accessibility correlates with expression of proximal genes. In normal human islets, clustering of significant peak-to-gene associations revealed distinct cell type-specific regulatory modules (**Fig. 5A, Suppl. Table 12**). Cluster 1 comprised regulatory elements whose accessibility positively correlated with expression of canonical beta cell marker genes in normal human islets. As a comparison, a similar cluster identified in insulinoma beta cells (cluster 5; **Fig. 5A**) exhibited accessible chromatin regions that were also associated with gene expression, serving as a control. Motif enrichment analysis of these clusters demonstrated that AP-1 motifs, particularly *JUNB*, were among the most significantly enriched motifs in cluster 1 but not in cluster 5, suggesting that AP-1-mediated enhancer activity is a defining feature of normal beta cell regulatory architecture and is disrupted in insulinomas (**Fig. 5B, Suppl. Table 13**).

**Figure 5:**
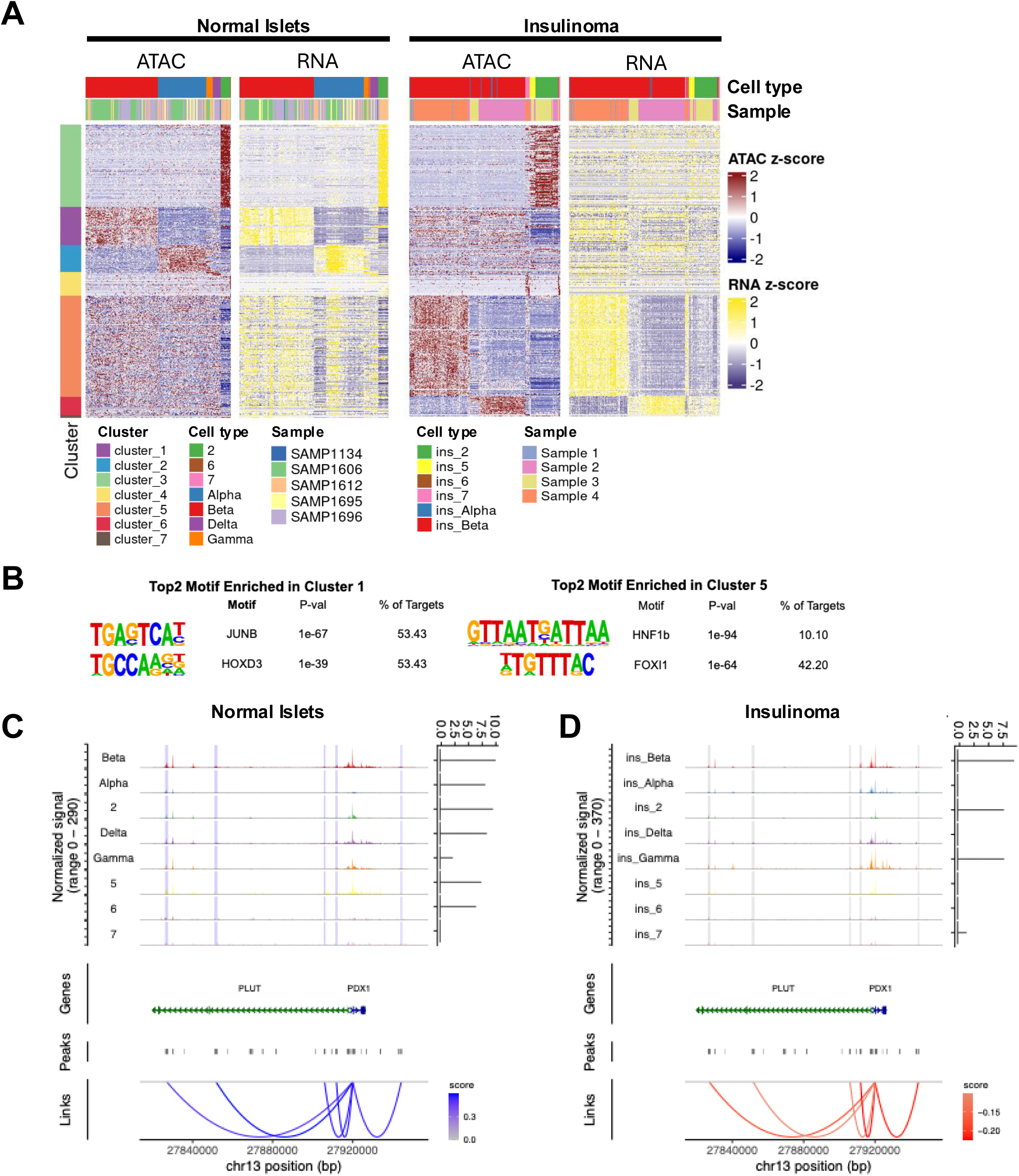
AP-1 Transcription Factors Regulate Beta Cell Gene Expression via Enhancer Elements. **A.** Heatmap of chromatin accessibility and gene expression for 8,577 unique significant peak-to-gene links across meta normal islet and insulinoma cells. Signal from 50 cells in each aggregated cell were summed. Significant peaks are determined through correlating gene expression and accessibility of ATAC-seq peaks within 2.5–250 kb of promoter. (Methods). Clusters 1–4 contain links identified in normal islet cells; clusters 5–7 contain links identified in insulinoma cells. **B.** Top 2 enriched de-novo transcription factor binding motifs in cluster 1 and cluster 5 from HOMER analysis. **C-D.** snATAC-seq genome browser tracks at PDX-1 locus for each cluster from normal islet (C) and insulinoma (D). Normalized and aggregated accessibility from snATAC-seq (top left). Normalized expression is presented on violin plot (top right). Highlighted regions represent the distal regulatory elements found in cluster1 in C that significantly correlate with PDX1 expression (blue) and insignificantly correlate with PDX1 expression (grey). Links from the significant distal regulatory elements to PDX1 TSS from cluster 1 (bottom). Note that these distal regulatory elements contain AP-1 binding motifs. Blue and red loops indicate positive and negative correlation with gene expression respectively.

To investigate locus-specific regulatory interactions, we examined peak-to-gene links surrounding the *PDX1* locus. In normal beta cells, distal enhancer elements containing AP-1 motifs exhibited strong positive correlations with *PDX1* expression (**Fig. 5C**). In contrast, although similar chromatin accessibility patterns were observed in insulinoma cells, these enhancer-promoter interactions lacked a positive correlation with gene expression (**Fig. 5D**), suggesting a functional decoupling of enhancer activity from transcriptional output of their associated gene. Comparable patterns were observed at additional beta cell marker gene loci, including *INS*, *MAFA*, and *PCSK1* (**Suppl. Fig. 10**).

Collectively, these findings support a model in which the AP-1 family transcription factors regulate beta cell identity through enhancer-mediated control of key lineage-specific genes, and that disruption of this regulatory axis contributes to the altered transcriptional landscape observed in insulinomas.

## Discussion

In this study, our goals were three-fold. First, although bulk RNA-seq and bulk proteomic approaches have suggested possible mechanisms underlying insulinoma molecular pathogenesis, we sought to define the exact cell types in which abnormal gene expression and chromatin regulation actually occurs in human insulinomas, and to compare these findings to normal human beta cells. Second, we hoped to link the well described association of epigenetic dysregulation in insulinomas^7^ to pathogenic mechanisms of action. And third, since some beta cells in benign insulinomas have the capacity to proliferate, expand their numbers and overproduce insulin, we hoped to find novel pathways in beta cells that could be exploited as novel targets for beta cell regenerative drug discovery for diabetes.

With regard to our first objective, we have clearly identified a recurring theme of reduced AP-1 transcription factor expression complimented by loss of transcription factor motif enrichment for the AP-1 family transcriptional regulators in accessible chromatin regions as a central and recurring feature among the beta cells of insulinomas. This novel observation may be attributable in part to the use of single cell/single nucleus approaches on beta cells and insulinomas. As an example, while JUN, JUND and FOS are expressed in beta cells, they are also highly expressed in ductal cells, acinar cells, endothelial cells (**Fig. 3D**) making changes in their expression in bulk samples problematic to interpret. Our single nucleus RNA-seq analysis has also clearly identified four transcriptionally distinct beta cells subtypes in insulinomas, one of which is characterized by a proliferative signature. Future studies will be necessary to further investigate this proliferating beta cell subtype which bears a great potential to discover novel targets/mechanisms for beta cell regenerative therapies. In addition, open chromatin profiling on a single cell level revealed six different subtypes of beta cells in insulinomas. Again, the detailed analysis delineating the differences of these beta cell subtypes and how they compare to normal islet beta cells will require additional work.

Importantly, our gene regulatory network analysis on insulinoma and normal islet datasets revealed that regulons driven by AP-1 family transcription factors were enriched in beta cells from normal islets but were substantially diminished in insulinoma beta cells. This, alongside chromatin accessibility and gene expression data, also provides supporting evidence that AP-1 transcriptional programs are required for maintaining beta cell identity. Indeed, silencing JUN in cadaveric human islets significantly reduced the expression of some of the canonical beta cell marker genes such as PDX-1, NKX6-1 and MAFA, whereas JUND silencing had minimal effect, indicating that JUN is the primary regulator of beta cell phenotype among AP-1 family members. Further, comparative single cell transcriptomic analysis of human islets from adult vs. prenatal donors showed that AP-1 family transcription factors, in particular JUN, JUNB and JUND, exhibit expression patterns that parallel established beta cell maturity markers. These data indicate that AP-1 family transcription factors are required for the acquisition and maintenance of a fully mature beta cell state and suggest that insulinoma beta cells adopt a more immature phenotype. Given that beta cell maturity is defined by glucose responsiveness threshold^21,22^, this immature state may contribute to the inappropriate insulin secretion observed in insulinomas, whereby beta cells secrete insulin at sub-physiologic glucose concentrations.

Our second key objective was to link events in normal and insulinoma beta cells in pathogenic terms. One obvious area of interest for AP-1-insulinoma pathogenesis was the well described relationship between JunD and MEN1. We and others previously demonstrated in the 1990’s and 2000’s that menin disruption causes beta cell expansion^23–29^, and that Menin and JunD are binding partners. In addition, it was shown that MEN1 is frequently mutated in human insulinomas and other PNETs and NENs, and that it serves as a member of the Trithorax or COMPASS epigenetic activating complex^5,11,23–29^. Agarwal and Marx showed that JunD binds to menin and is required for its repressive activity^23,24,30^. Huang et al demonstrated that MLL1 and MLL2 both bind to, and compete with one another for binding to, menin^31^. Our studies make the novel point that, like MEN1 mutations, AP-1 dysregulation and AP-1 chromatin inaccessibility, particularly at enhancers, are two very specific features of human insulinoma biology. Understanding why certain chromatin regions/enhancers become inaccessible, why simultaneous AP-1 expression is decreased, and defining the key target genes for AP-1s, menin, and MLLs in human beta cells and insulinomas are critical next steps. Stated more succinctly, why do multiple mutations and variants in different epigenetic regulatory genes all converge specifically on AP-1 genes and their enhancer binding sites, and how does this “generic” transcription factor family lead to a very specific and reproducible beta cell phenotype? Addressing these important questions will require additional studies.

In addition to epigenetic regulatory gene variants, altered DNA methylation has been observed repeatedly in insulinomas. For example, Chan et al. and our group^7,32^ have both reported genome-wide DNA methylome abnormalities in human insulinomas. In particular, abnormal DNA methylation patterns at the imprinted *CDKN1C* locus leads to silencing of the *CDKN1C* product, the cell cycle inhibitor p57^KIP2^, and this is permissive for human beta cell proliferation. But how and why this locus is targeted for altered methylation is unknown. We speculate that AP-1 expression and/or chromatin access may play a role here.

Although the JunD-Menin axis in insulinomas was discovered more than 25 years ago, very little is known about the role of the AP-1 transcription factors in human insulinoma. More specifically, given the prior reports suggesting that JunD is a menin binding partner^23^, we were surprised that silencing JunD had little effect. AP-1s may bind to their target genomic motifs as homodimers (e.g., Jun-Jun) or heterodimers e.g., JunD-Fos. They may also form heterodimers with other families of transcription factors, such as the CREB superfamily^33^, the ATF family and/or the NFAT family^34^. Our gene expression data make some of these unlikely (e.g., FosL1, FosL2, ATF1,3,5, BATF1-3, CREBs 5, 3L4, 3L1, and NFATs 1,2 are all negligibly expressed in beta cells), but others are abundant and are reasonable candidates for AP-1 transcriptional partners. Thus, key unanswered mechanistic questions are “where in the beta cell genome AP-1s bind”, “what specific beta cell-relevant targets do they regulate”, “do they serve as canonical activators or as repressors as occurs in partnership with JunD and menin”, and “do they act as heterodimers or homodimers, and if so, what are the key AP-1 heteromeric partners.”

The UMAPs in Suppl. Fig.5, make it is clear that each for the four insulinomas has its own unique components of beta cell subtypes. This is expected, since it is likely that they all had different driver mutations, downstream gene expression patterns and accessible chromatin alterations. As an example, one might expect the insulinoma beta cells in a MEN1 mutant insulinoma to cluster separately from those of a YY-1 mutant insulinoma, or a KDM6A mutant insulinoma. Defining the key AP-1 targets lost or gained in insulinomas is an important future goal.

Our third key objective, building off the observation that insulinoma beta cells very effectively proliferate, produce insulin, lower blood glucose, and increase beta cell mass, is to identify regulatory molecules within beta cells that might serve as targets for enhancing human beta cel numbers and function for people with diabetes. As an example, in our earlier report on insulinomas, we found that insulinomas displayed marked alterations in DYRK1A RNA splicing. We and others have shown that small molecule inhibitors of the kinase DYRK1A, enhance human beta cell mass and function, and reverse diabetes. As another example, small molecule interference with TGFβ superfamily signaling enhances human beta cell proliferation^13,14^. In the current study, we demonstrated that AP-1 family transcription factors, in particular JUN is critical to regulate beta cell proliferation and differentiation. Further, analysis of published single cell datasets from human islets from adult and prenatal donors showed that JUN, JUNB and JUND mimicked the expression pattern of well-established beta cell marker genes, such as PDX-1, NKX6-1 and MAFA (**Fig. 4G**), suggesting a role for AP-1 members in beta cell maturation and functional differentiation. This idea is further supported by Alvarez-Dominguez et al. where they demonstrated that DNA binding motifs for AP-1 family members, in particular JUNB and FOS, are highly enriched in mature primary beta cells when compared to stem cell-derived islets^35^. As another example, a small molecule inhibitor of menin, BMF-219 or Icovamenib, is currently in clinical trials in people with diabetes as a beta cell regenerative therapeutic (Clinicaltrials.gov ID: NCT06152042). Thus, learning more about precisely how AP-1 factors regulate human insulinoma cell proliferation and maturation may provide additional targets for human beta cell regenerative drug discovery.

### Limitations

This study has limitations. As noted above, defining specifically how AP-1 binding sites become inaccessible and how AP-1 expression is coordinately reduced will require further study. Similarly, how AP-1 members affect activation of specific beta cell relevant genes requires deeper exploration. If and how AP-1s might also regulate human embryonic stem cell therapeutic programs, and whether drugs that activate AP-1s in human beta cells might be of therapeutic value provide opportunities for deeper investigation. Another limitation is the absence of whole genome or whole exome sequencing on the insulinoma samples we have studied. Going forward, extending these kinds of single cell studies to additional, larger numbers of human insulinomas coupled with whole exome sequencing and AP-1 family ChIP-seq studies will be critical to clarifying gene target interactions and to determine if what appears to be a homogeneous AP-1 insulinoma relationship will extend to larger sample sets.

## Methods

### Human Pancreatic Islets

Isolated de-identified human pancreatic islets from otherwise normal organ donors were provided by the NIH Integrated Islet Distribution Program (IIDP, https://iidp.coh.org), Prodo Laboratories, The Alberta Diabetes Institute, and the Transplant Surgery Department, University of Chicago. Details and demographics of the donors and islet preparations are provided in Suppl. Table 14. Donor ages ranged from 26 to 66 years old. The mean age (±SEM) was 48 ± .087 years; mean BMI (±SEM) was 26.67 ± 0.4 kg/m2 (range 20.7-39.2).

### Human Insulinomas

Four human insulinomas were collected from subjects who provided informed written consent and were deposited in the ISMMS Biorepository. Patient samples were de-identified through the Biorepository and Pathology Core at the ISMMS (IRB-HSM-00145) or the providing institutions. Clinical details of the patients with insulinoma are provided in Supplementary Table 15.

### Bulk RNAseq Differential Gene Expression Analysis

Bulk RNA-seq read counts from 22 sorted beta-cell samples and 34 insulinoma samples (generated internally^7^ or obtained from published datasets^5,10,36,37^) were analyzed in R (v4.2.2) using edgeR (v4.4.2). Genes were retained if they had ≥10 reads in ≥70% of samples in each group. Library composition bias was corrected using TMM normalization. Differential expression was tested with an edgeR negative binomial generalized linear model including group (beta cells vs insulinomas), sex, and sequencing source (internal vs literature-derived) as covariates. Dispersion was estimated using robust empirical Bayes procedures (estimateDisp followed by tagwise dispersion estimation), and gene-wise differential expression for the group term was assessed using a likelihood ratio test (LRT). Results are reported as log2 fold change (logFC) with Benjamini–Hochberg FDR correction; significance was defined as FDR < 0.05.

### Bulk ATAC-seq Experiments

Nuclei were isolated from FACS-sorted beta cells and Tn5 transposition was performed following a published protocol^38^. 50,000 nuclei were used from each FACS-sorted beta cell sample. Nuclei were also isolated from flash frozen insulinomas and Tn5 transposition was performed according to the OMNI-ATAC protocol^39^. 20-30mg flash-frozen tissue was used from each insulinoma. ATAC-seq libraries were prepared using Nextera Library Prep kit from Illumina (FC-121-1030). Samples were sequenced on Illumina NextSeq 500 instruments.

### H3K27Ac ChIP-seq Experiments

H3K27Ac profiling on 3 FACS-sorted beta cells and 4 insulinomas was performed. FACS-sorted beta cells were collected after 4 days of adenoviral infection of Zs-Green that is expressed under the control of *INS* promoter^7^. Chromatin was prepared and the ChIP was performed on FACS sorted beta cells and insulinomas following ChIPmentation protocol^40^ which is a fast and robust method for low input ChIP-seq. ChIP-seq libraries were prepared using Nextera Library Prep kit and Nextera adapters from Illumina (#15032354). Samples were sequenced on Illumina NovaSeq 500 instruments.

### Bulk ATAC-seq Processing

Bulk ATAC-seq was performed on n = 4 human insulinomas and n = 3 FACS-sorted beta cells. Briefly, read quality was assessed using FastQC (v0.11.8)^41^. Adapter sequences were trimmed using Trim Galore! (v0.6.6)^42^, and reads were aligned to the GRCh37.p13 assembly of the human genome using Bowtie2 (v2.2.8)^43^. Reads aligned to mitochondrial DNA and those with mapping quality Q < 20 were removed. Picard (v2.2.4) was used to remove duplicated reads^44^. Post-filtering BAM files were merged using SAMtools (v1.11) merge^45,46^, followed by peak calling using MACS2 (v2.1.0) with parameters ‘--nomodel --nolambda --keepdup all --slocal 10000^47,48^. Peaks overlapping ENCODE hg19 blacklisted regions were removed. Quantification of reads in peaks was performed using BEDTools multicov (v2.29.2)^49^. Peaks with fewer than 200 total reads across all samples were filtered as inactive peaks. Differential peak analysis was performed using DESeq2 (v1.30.1); peaks with adjusted P-value < 0.01 and absolute log2 fold change ≥ 2 were considered differentially accessible^50^. Coverage in BAM files was normalized by the median-of-ratios scaling factor calculated from reads aligned to called peaks in DESeq2 and converted to bigWig files using deepTools bamCoverage (v3.2.1)^51^. deepTools computeMatrix and plotHeatmap functions were used to generate heatmaps.

### Motif Analysis of Bulk ATAC-seq Data

Motif enrichment analysis was performed using Analysis of Motif Enrichment (AME) from the MEME Suite^52^, implemented via the R package memes (v1.2.5)^53^. Regions for motif analysis were extended 100bp upstream and downstream from peak summit, and the underlying DNA sequences were extracted from the hg19 reference genome using the BSgenome.Hsapiens.UCSC.hg19 R package (v1.4.3). AME was run using the JASPAR 2022 CORE vertebrates non-redundant position frequency matrix database ^54^ against a background set of 10,000 randomly subsampled non-differentially accessible (static) ATAC-seq peaks that did not overlap with detectable H3K27Ac signal. To focus on biologically relevant candidates, motif dimers were excluded and only TFs with detectable expression in the bulk RNA-seq dataset were retained. Results were visualized as lollipop plot in which the size of each dot represents the significance of enrichment and the y-axis shows the percentage of peaks containing the specific motif. The top 10 motifs from each test set of peaks were plotted. The lollipop plot was generated using ggplot2^55^.

### TOBIAS Analysis of Bulk ATAC-seq

Footprinting analysis was performed using TOBIAS (v0.13.2) on aggregated BAM files per condition^56^. The ATACorrect function was used to correct Tn5 insertion bias, and ScoreBigwig was used to calculate TF-binding scores in peak regions. Differential binding fold change and P-values for each TF were determined using the BINDetect function, and selected TF results were visualized using PlotAggregate. Volcano plots were generated using ggplot2 to visualize differential binding between insulinoma and sorted beta cells at all ATAC-seq peaks and at ATAC-seq peaks overlapping with beta cell–specific, insulinoma-specific, or shared enhancers. As with the AME analysis, motif dimers were excluded and only TFs with detectable expression in bulk RNA-seq were retained.

### H3K27Ac ChIP-seq Analysis

In total 7 libraries were sequenced (5 samples from batch1 as 74 bp single-end, 2 samples from batch2 were sequenced as 74 bp paired-end). Read quality was assessed using FastQC. Adapter sequences were trimmed using Trim Galore!, and reads were aligned to the GRCh37.p13 assembly of the human genome using Bowtie (v1.3.0) with parameters -k 1 -m 1 --best -n 2 -l 70 --chunkmbs 50^57^. SAMtools (v1.11) was used to generate BAM files and filter out low-quality (Q < 20) and mitochondrial reads. Picard (v2.2.4) was used to remove duplicate reads. Coverage tracks (bigWig files) were generated from filtered BAM files using deepTools (v3.5.1) bamCoverage with parameters ‘--normalizeUsing RPKM --binsize 10 --extendReads 200’. BAM files from sample H218-Zsplus-Dia-K27Ac_S9 (Beta_2) and sample 12728-K27Ac_S15 (INS_1) were merged using SAMtools merge, and significant binding peaks of the merged datasets were determined using MACS2 (v2.1.0) with the input file as control and -q 0.1. Peaks located in ENCODE hg19 blacklisted regions were excluded. Nearby peaks (<12.5 kb apart) in the filtered peak set were stitched together, and peaks overlapping promoters (± 2kb from TSS) were excluded using ROSE (v1.0)^58,59^. DiffBind (v3.4.11)^60^ was applied to identify the H3K27Ac regions that increases or decreases binding between insulinoma and normal islet and overlap with ATAC-seq peaks.

To delineate cis-regulatory regions associated with beta cells, insulinoma, or both, we integrated ATAC-seq with H3K27Ac ChIP-seq data by classifying each ATAC-seq peak along two features, differential accessibility status and H3K27Ac enrichment status at that locus (overlap with a beta-cell enriched, insulinoma enriched, or static H3K27Ac peak, or absence of any called H3K27Ac peak). This two-feature classification initially yielded eight categories, which were merged into four major clusters (the fourth cluster which missing H3K27Ac signal is not shown) based on the concordance of chromatin accessibility and H3K27Ac signal: beta-cell-associated cis-regulatory regions (c1,c3,c6), insulinoma-associated cis-regulatory regions (c2), and shared cis-regulatory regions (c4, c5). deepTools computeMatrix and plotHeatmap functions were used to generate heatmap to visualize the signal in each cluster across samples.

### Functional Enrichment Analysis

To assess the biological functions associated with each peak cluster, genomic region enrichment analysis was performed using the rGREAT R package (v2.4.0)^61^. ATAC-seq regions associated with Beta Cell, Insulinoma or shared (**Fig1A**) were used as input. Peaks were assigned to genes using the “basal plus extension” rule, with a basal regulatory domain of 5 kb upstream and 1 kb downstream of each TSS, extended up to 1 Mb until the neighboring gene’s basal domain was reached. TSS annotations were obtained from the GREAT hg19 reference^62^. To restrict the analysis to a biologically relevant background, the gene universe was defined as all genes detected in the bulk RNA-seq datasets. Enrichment was computed against the KEGG gene set collection from MSigDB (v2025.1)^63^. For visualization, the top 5 enriched pathways per cluster (by adjusted P-value) were selected, and the union of these terms was displayed across all three clusters in dot plots.

### Single-Nucleus RNA-seq: Insulinoma Sample Quality Control and Preprocessing

FASTQ files were aligned to the GRCh38 human genome reference (GRCh38-2020-A-2.0.0), filtered, barcoded, and UMI-counted using Cell Ranger (v7.2.0, 10x Genomics)^64^. Empty droplets and ambient RNA were removed from count matrices using the remove-background function from CellBender (v0.2.2)^65^. Each dataset was then filtered to retain cells with ≥1,000 UMIs, ≥400 expressed genes, and <10% of reads aligned to the mitochondrial genome. UMI counts were normalized so that each cell had a total of 10,000 UMIs across all genes and then log-transformed with a pseudocount of 1 using the LogNormalize function in the Seurat package (v4.0.3)^66^. The top 2,000 most highly variable genes were identified using the vst selection method of FindVariableFeatures, and counts were scaled using ScaleData. Principal component analysis (PCA) was performed using the top 2,000 highly variable features (RunPCA), and the top 30 principal components were used in downstream analysis.

### Single-Nucleus ATAC-seq: Insulinoma Sample Quality Control and Preprocessing

FASTQ files from three insulinoma samples were processed using Cell Ranger ATAC (v2.1.0, 10x Genomics) with the GRCh38 reference genome (GRCh38-2020-A-2.0.0). Peaks were called using the CallPeaks (MACS2) function in Signac (v1.3.0)^67^. Cells were filtered based on the following criteria: fragments in peaks >3,000 and <30,000, >20% reads in peaks, blacklist ratio <0.05, nucleosome signal <4, and TSS enrichment score >3. The blacklist ratio is defined as the fraction of reads aligned to blacklisted regions relative to total reads. Nucleosome signal was calculated using NucleosomeSignal in Signac, and TSS enrichment score was calculated using TSSEnrichment. Latent semantic indexing (LSI) was applied to the top 95% most variable peaks identified using FindTopFeatures, combining term frequency–inverse document frequency (TF-IDF) normalization with singular value decomposition (SVD) for dimensionality reduction. The first LSI component was excluded because of its correlation with sequencing depth. De novo clustering (resolution 0.2) was performed with FindNeighbors and FindClusters in Signac.

### Single-Nucleus Multiome (ATAC + Gene Expression) Quality Control and Preprocessing

FASTQ files from one insulinoma single-nucleus multiome sample were aligned, filtered, barcoded, and UMI-counted using Cell Ranger ARC (v2.0.0, 10x Genomics) with the GRCh38 reference genome (GRCh38-2020-A-2.0.0). Sequenced FASTQ files from the published normal islet dataset (GSE200044)^68^ were retrieved from GEO and processed using Cell Ranger ARC (v2.0.2) with the same reference genome. Both datasets followed the same post-alignment quality control and preprocessing steps as described in the snRNA-seq insulinoma preprocessing section. Briefly, the snRNA-seq portion of each dataset was filtered to retain cells with ≥1,000 UMIs, ≥400 expressed genes, and <10% mitochondrial reads. UMI counts were normalized, and the top 2,000 most highly variable genes were selected for PCA. For the snATAC-seq portion, peaks were called on individual samples, and cells were filtered using the same criteria as for the insulinoma snATAC-seq samples (fragments in peaks >3,000 and <30,000, reads in peaks >20%, blacklist ratio <0.05, nucleosome signal <4, TSS enrichment score >3). LSI was applied to the top 95% most variable peaks using TF-IDF normalization and SVD. The first LSI component was excluded owing to its correlation with sequencing depth.

### Integration of snRNA-seq Samples

The two snRNA-seq insulinoma samples were integrated with the RNA portion of the insulinoma multiome sample using Seurat’s integration workflow. The top 2,000 integration features were selected using SelectIntegrationFeatures, and SCTransform-normalized data were prepared using PrepSCTIntegration. Integration anchors were identified using FindIntegrationAnchors, and samples were integrated using IntegrateData. ScaleData and RunPCA were applied for scaling and PCA on the integrated dataset. K-nearest neighbor (KNN) graphs were constructed using FindNeighbors based on the top 30 principal components, and UMAP embeddings were computed using RunUMAP. The Louvain algorithm was used to cluster cells based on expression similarity via FindClusters using a resolution of 0.2. Clusters were annotated based on the expression of canonical pancreatic islet cell type markers. The proportion of each sample within each annotated cell type was calculated as the number of cells from a given sample in a cell type divided by the total number of cells of that cell type in the integrated dataset. The preprocessed snRNA-seq portion of the six normal islet multiome samples (GSE200044) was also integrated using the same workflow. Cells were annotated using the original labels from GSE200044.

### Integration of snATAC-seq Samples

The three snATAC-seq insulinoma samples were integrated with the ATAC portion of the insulinoma multiome sample by first unifying peaks called from individual samples. Aligned reads at the unified peak set were then quantified across cells from all samples to construct a peak-by-cell count matrix, which was used to create a Seurat object. TF-IDF normalization was applied, followed by SVD for dimensionality reduction. Harmony^69^ was used to correct for batch effects. De novo clustering (resolution 0.2) was performed using FindNeighbors and FindClusters in Signac. De novo clusters were annotated using aggregated accessibility at canonical pancreatic islet marker genes. Following annotation, peaks were re-called on each individual annotated cluster using MACS2, and cluster-specific peaks were defined as those uniquely identified in a given cluster with no overlap with peaks called in any other cluster, enabling assessment of chromatin accessibility differences at the cluster level. Peaks were annotated to genomic features using annotatePeak from ChIPseeker (v1.36.0)^70^ R package, and the distribution across promoters, introns, exons, UTRs, and intergenic regions was visualized as a pie chart. Motif position frequency matrices were retrieved from the JASPAR 2022 database and added to the integrated snATAC-seq data using AddMotifs in Signac. ChromVAR motif analysis was performed using RunChromVAR^71^, and FindAllMarkers from Seurat was used to apply the Wilcoxon rank-sum test to identify TF motifs significantly active or inactive in each cluster relative to all other clusters (adjusted p-value <0.01 and absolute log2 foldchange > 0.25). The scaled chromVAR deviation scores of the top 5 TFs from each cluster were displayed as a heatmap. Selected TFs were visualized on UMAP using FeaturePlot. The preprocessed snATAC-seq portion of the six normal islet multiome samples (GSE200044) was also integrated using the same workflow, and cells were annotated using the original labels from GSE200044.

### Projecting Insulinoma Samples onto Normal Islet Reference

To identify the most similar normal islet cell type for each insulinoma cell, we performed reference mapping using Seurat’s MapQuery workflow. Transfer anchors were identified using FindTransferAnchors, with the normal islet integrated dataset as the reference and the insulinoma integrated dataset as the query. For RNA, the first 30 PCA dimensions were used as the reference reduction with SCTransform normalization. For ATAC, LSI dimensions 2–30 were used as the reference reduction. Projection was performed using MapQuery, and projection results were visualized on the reference UMAP space. Subsequently, all insulinoma and normal islet samples were integrated using the same workflow described for insulinoma samples alone to enable downstream differential analyses. Motif position frequency matrices from JASPAR 2022 were added to the integrated snATAC-seq data. chromVAR motif analysis was performed using RunChromVAR, and FindMarkers from Seurat was used to apply the Wilcoxon rank-sum test to identify TF motifs significantly enriched or depleted between insulinoma and normal islet cells within each cell type (adjusted p-value <0.01 and absolute log2 foldchange > 0.25).

### SCENIC+ Enhancer-Driven Gene Regulatory Network Inference

The SCENIC+ (v1.0a1) pipeline^72^ was implemented as a Snakemake workflow^73^ to infer enhancer-driven gene regulatory networks (eGRNs) from integrated single-nucleus (sn) RNA-seq and ATAC-seq data (10 samples for ATAC: 6 normal islet and 4 insulinoma; 9 samples for RNA: 6 normal islet and 3 insulinoma). Both snRNA-seq and snATAC-seq data of insulinoma and normal islet were batch corrected using harmony. Chromatin accessibility count matrices were exported from Signac and used to construct a cisTopic object using pycisTopic (v2.0a0)^74^. LDA topic modeling was performed using collapsed Gibbs sampling (500 iterations)^75^, and the optimal model (32 topics) was selected based on established quality metrics. The cell-by-gene and cell-by-peak matrices were then submitted to the SCENIC+ pipeline, which generated paired snRNA-seq and snATAC-seq pseudo-multiome metacells from the input data. TF motif enrichment was scored using cisTarget and the Differential Enrichment Method (DEM) with hg38 v10 non-redundant clustered motif databases, within a search space of ±100 kb from each transcription start site. TFs, their target regulatory regions, and target genes were identified across cell types. We focused on direct eRegulons, and the most robust regulons were prioritized based on SCENIC+ recommendations (activators, +/+; repressors, −/−). eRegulon specificity scores (RSS) were computed to identify cell-type-specific regulons. The activator eRegulons from the top 50 most variable eRegulons, ranked by variance of gene-based AUC scores across cells and restricted to classic activator and repressor configurations, were visualized as a heatmap-dotplot, where color intensity represents mean gene-based AUC and dot size represents mean region-based AUC per cell type cluster.

### Matching snATAC-seq to snRNA-seq and Distal Regulatory Element-to-Gene Linkage

To integrate chromatin accessibility with gene expression data, we employed a strategy adapted from Granja et al.^76^. Cicero^77^ was run separately on non-diabetic and insulinoma snATAC-seq cells to generate k-nearest-neighbor (KNN) metacell groupings (k = 50), compute gene activity scores, and identify co-accessible peak links within each condition. Co-accessible links from both conditions were then merged for downstream comparison. For the six non-diabetic donors, paired snRNA-seq and snATAC-seq data were available from the same multiome experiment, so cells from snATAC-seq and from snRNA-seq were directly matched. For the insulinoma samples, which lacked paired multiome data, gene activity scores were then used to align snATAC-seq cells to their most similar snRNA-seq counterparts via canonical correlation analysis (CCA) in Seurat. Briefly, Seurat objects were created from the Cicero gene activity score matrix and the snRNA-seq count matrix, each normalized and scaled independently. The union of the top 2,500 most variable features from both modalities was used to run CCA with 20 canonical correlations. For each snATAC-seq insulinoma cell, the nearest snRNA-seq insulinoma cell was identified by minimizing Euclidean distance in the CCA-aligned embedding. Matched pairs with a CCA correlation below 0.4 were excluded.

Using the initial KNN groupings from Cicero, aggregated snATAC-seq accessibility and matched snRNA-seq expression matrices were constructed at the metacell level for both insulinoma and normal islet. Metacell cell type identity was assigned by majority vote among KNN neighbors. Due to the low abundance of gamma and delta cells in the insulinoma samples, insulinoma version of gamma and delta cells did not form a plurality in any metacell neighborhood and were therefore not represented as distinct cell type clusters in the peak-to-gene linkage analysis. To identify peak-to-gene links, each gene’s genomic range was resized to its transcription start site and extended to a ±250 kb window. All ATAC-seq peaks overlapping this window were identified as putative peak-to-gene pairs using findOverlaps from the GenomicRanges package (v1.52.0)^78^. Peak-to-gene correlations were computed separately for non-diabetic and insulinoma metacells. To establish statistical significance, a null correlation distribution was computed from trans correlations (correlating 1,000 peaks on different chromosomes with each gene) and used to derive z-scores, P-values, and Benjamini–Hochberg-adjusted FDR values for each putative link. Significant peak-to-gene links were defined as those with Pearson correlation ≥ 0.5, FDR < 0.1, and a minimum distance of 2.5 kb from the TSS. Links were classified as non-diabetic-specific, insulinoma-specific, or shared based on whether they met significance thresholds in one or both conditions.

### Motif Scanning and Enrichment Analysis of Distal Regulatory Element-to-Gene-Linked Regions

To identify transcription factor motifs enriched in peak-to-gene-linked regulatory regions, we performed motif enrichment analysis using HOMER findMotifsGenome.pl (v5.1)^79^. HOMER was run on peaks from each of the seven peak-to-gene link clusters identified by the distal regulatory element-to-gene linkage analysis. Peaks were provided in BED format with their original genomic coordinates (-size given) against the hg38 genome.

For targeted AP-1 motif scanning, FIMO (v1.2.5)^80^ from the MEME Suite was applied to scan the sequences underlying all significant peak-to-gene links. Peak sequences were extracted from the hg38 genome (BSgenome.Hsapiens.UCSC.hg38) and scanned against the JASPAR 2022 human motif database using a significance threshold of P < 1 × 10⁻⁴. Motif hits were intersected with peak-to-gene link metadata to identify peaks harboring AP-1 family motifs (JUN, JUND, JUNB, FOS, FOSL1, FOSL2, ATF, and Fra family members). Chromatin accessibility coverage tracks at beta cell marker gene loci were generated using Signac’s CoveragePlot, with peaks containing AP-1 motifs highlighted. Peaks were color-coded by whether they were significantly linked in non-diabetic samples only, insulinoma samples only, or both conditions.

### Bulk RNA-seq differential gene expression analysis

Bulk RNA-seq read counts from 22 sorted beta-cell samples and 34 insulinoma samples (generated internally^7^ or obtained from published datasets^5,10,36,37^) were analyzed in R (v4.2.2) using edgeR (v4.4.2). Genes were retained if they had ≥10 reads in ≥70% of samples in each group. Library composition bias was corrected using TMM normalization. Differential expression was tested with an edgeR negative binomial generalized linear model including group (beta cells vs insulinomas), sex, and sequencing source (internal vs literature-derived) as covariates. Dispersion was estimated using robust empirical Bayes procedures (estimateDisp followed by tagwise dispersion estimation), and gene-wise differential expression for the group term was assessed using a likelihood ratio test (LRT). Results are reported as log2 fold change (logFC) with Benjamini–Hochberg FDR correction; significance was defined as FDR < 0.05.

### scRNA-seq dataset merging and integration for the comparison of single cell datasets from adult and prenatal donors

Single-cell RNA-seq data were compiled from multiple publicly available human pancreatic islet studies (Supplementary Table 16) and standardized into Seurat (v5.0.0) objects in R (v4.4.2) with harmonized metadata and a unified gene identifier space. For datasets not distributed in 10x format, count matrices were reconstructed into standard 10x-style files and imported; when provided as Seurat objects, raw counts were extracted from the RNA assay and reformatted consistently. Study-specific metadata were curated to capture subject-level covariates.

Gene identifiers were harmonized by mapping gene symbols to Ensembl IDs using hg38 annotation (hg19 as fallback when required), with symbol sanitation, duplicate checks, and recovery of legacy symbols using GeneSymbolThesarus() (previous-symbol lookup). A final feature table (Ensembl ID, gene symbol) was generated after excluding problematic mappings.

Each dataset underwent standard Seurat preprocessing (normalization, variable feature selection, scaling, PCA/UMAP). Doublets were identified with DoubletFinder (v2.0.6) using dataset-specific parameter selection and removed prior to downstream analysis. Cell-level metadata were standardized across studies (e.g., study, subject, sex, age, nCount_RNA, nFeature_RNA, doublet status).

Study objects were merged, genes were retained if detected in ≥10 cells, and QC filtering was applied, including mitochondrial fraction calculation and thresholds for library size, detected features, and mitochondrial content; all-zero cells were removed. The merged dataset was reprocessed with PCA/UMAP and batch corrected with Harmony (v1.2.3) using study as the batch variable, followed by neighbor graph construction and clustering. Integration quality was assessed by comparing embeddings pre/post batch correction and by inspecting endocrine marker expression (e.g., INS, GCG, SST).

Cell-type labels were assigned with Azimuth (v0.5.0) using the pancreasref reference, and annotation quality was evaluated using prediction scores. Marker genes for Azimuth-predicted populations were identified using FindAllMarkers() with standard thresholds, and marker expression/cell-type distributions were visualized using standard Seurat plotting functions, including DotPlot().

### Human Islet Dispersion and Virus Infection

Islets were centrifuged at 1500 rpm for 5 min, washed twice in phosphate-buffered saline (PBS), re-suspended in 1 ml of Accutase and incubated for 10 min at 37°C. During digestion, the islets were dispersed by gentle pipetting up and down every 5 min for 10 sec. Complete RPMI medium containing 11 mmol/L glucose, 1% penicillin/streptomycin with 10% fetal bovine serum (FBS) was then added to stop the digestion. Dispersed cells were then centrifuged for 5 min at 1500rpm, the supernatants removed, the pellets re-suspended in complete medium, and the cells then plated on coverslips with 30 µl of cell suspension per coverslip. Poly-D-Lysine/Laminin-treated cover slides or chamber slides were used. Cells were allowed to attach for 2 hr at 37°C or were transduced with adenovirus for 2 hr. After 2 hr, 500 µl complete RPMI medium was added in each well to terminate adenoviral transduction. Cells were cultured for 96 hours. Detailed protocols are provided in reference^81^.

### Adenoviruses and Transduction

For adenoviral silencing, four target shRNA sequences for each target gene were designed using the Thermo Fisher RNAidesigner online tool. Target sequences were cloned into block-iT U6 RNAi entry vector (K494500, Invitrogen), and silencing efficiency was evaluated in HEK293 cells. The most effective target sequences were used to generate Ad-shRNAs in the Block-iT adenoviral RNAi vector (K494100, Invitrogen). Silencing sequences were as follows: shJUN: GCAAACCTCAGCAACTTCAAC, shJUND: GGAGGATTTACACAAGCAGAA.

### qRT-PCR Methods

RNA was isolated and quantitative RT-PCR was performed as described previously^82^ (46). Gene expression in dispersed islets was analyzed by real-time PCR performed on a QuantStudio System. Sequence information for primers is available in our previous publications and listed in Suppl. Table 17^13,83–85^.

### Statistics

Statistical analyses for qRT-PCR experiments were performed using 2-tailed Student’s paired t-test as described in the Figure Legends. P-values less than 0.05 were considered to be significant.

## Data Availability

All sequencing data are available on GEO with accession number GSE327169.

## Acknowledgments

The authors wish to thank Bonnie and Joel Bergstein, Lonnie and Thomas Schwartz, and Martha and Fred Farkouh families for their constant support of this research. We also thank the NIDDK-supported Human Islet and Adenovirus Core (HIAC) of the Einstein-Sinai Diabetes Research Center (ES-DRC), and the NIDDK Integrated Islet Distribution Program (IIDP), Prodo Laboratories, Patrick MacDonald at the Alberta Diabetes Institute Islet Core at the University of Alberta in Edmonton (www.bcell.org/adi-isletcore) with the assistance of the Human Organ Procurement and Exchange (HOPE) program, Trillium Gift of Life Network (TGLN), and other Canadian organ procurement organizations. We also thank Dr. Tatsuya Kin at the University of Alberta, Edmonton, Alberta, Canada, Dr. Piotr Witkowski at the University of Chicago, Chicago, Illinois, and Dr. Fouad Kandeel at the City of Hope Medical Center, Duarte, California, for providing human organ donor islets. We also thank Drs Steve Libutti, William Inabnet, Rajesh V. Thakker, Hyunsuk Suh, Mark Stevenson and Ira Goldberg for providing insulinoma surgical specimens. We also thank Alan Soto and Yayoi Kinoshita from Biorepository and Pathology CoRE, Rachel Chen and Dr. Zhihong Chen from Human Immune Monitoring Center and to Sanjana Shroff, Drs. Kristin Beaumont and Robert Sebra from Genomics Core Facility at the Icahn School of Medicine at Mount Sinai. This work was supported in part by the Bioinformatics for Next Generation Sequencing (BiNGS) shared resource facility within the Tisch Cancer Institute at the Icahn School of Medicine at Mount Sinai, which is partially supported by NIH grant P30CA196521. This work was also supported in part through the computational resources and staff expertise provided by Scientific Computing at the Icahn School of Medicine at Mount Sinai and supported by the Clinical and Translational Science Awards (CTSA) grant UL1TR004419 from the National Center for Advancing Translational Sciences. Research reported in this paper was supported by the Office of Research Infrastructure of the National Institutes of Health under award number S10OD026880 and S10OD030463. This work was supported by NIH grants P30DK020541, R01DK116873, R01DK116904, R01DK125285, R01DK105015, R01DK129196, R01DK139631.

## Author Contributions

E.K., D.H. and A.F.S. conceived of the studies. E.K., P.W., H.L. and D.H. performed experiments. X.W., L.L., E.K, D.H. and A.F.S. analyzed data. E.K. and A.F.S. wrote the manuscript with comments from X.W and D.H.

## Guarantors

E.K, D.H. X.W. L.L. and A.F.S. guarantee the data in this manuscript.

---

**Supplementary Figure 1.**
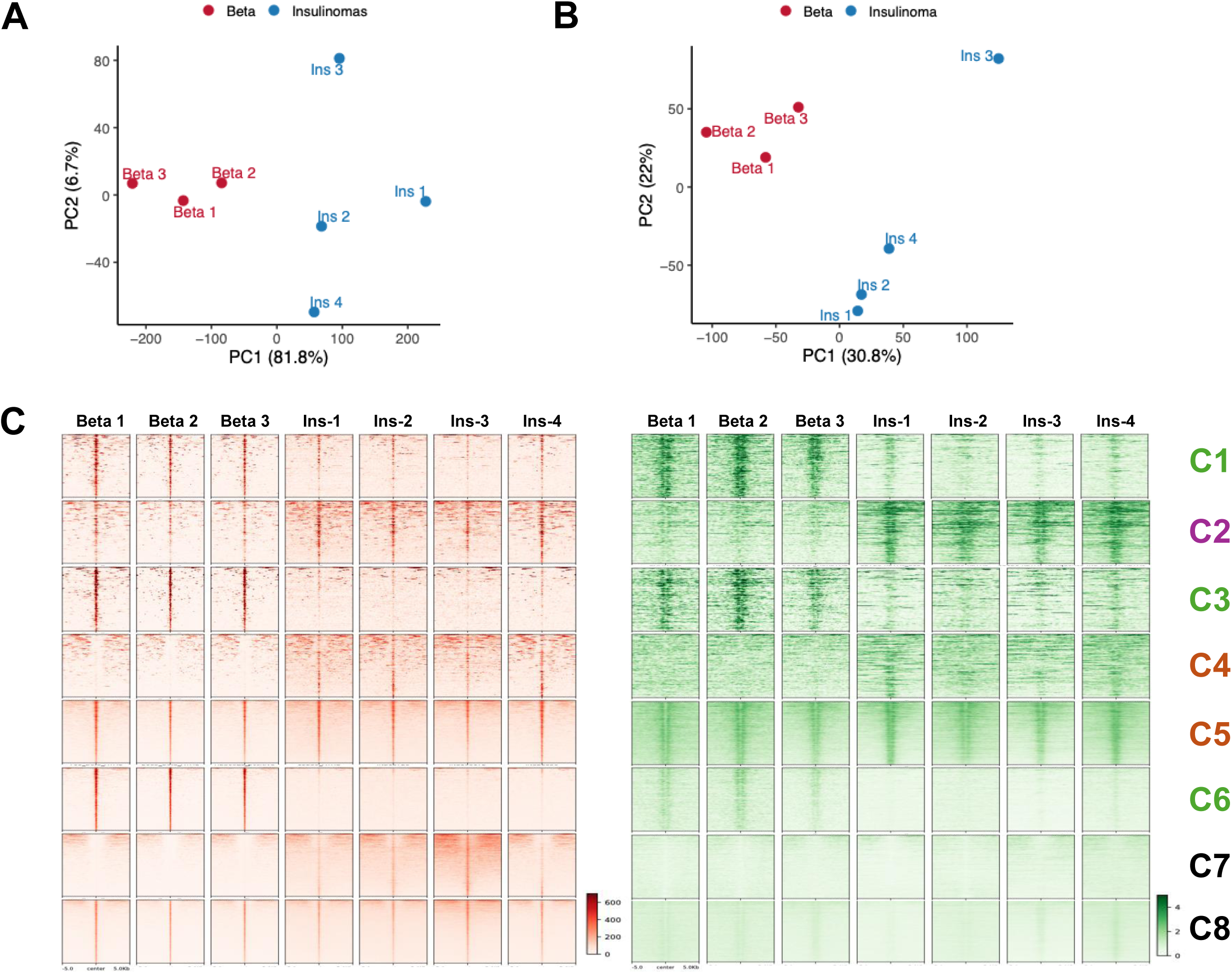
**A.** Principal component analysis (PCA) of normalized bulk ATAC-seq data, performed on the top 10,000 most variable peaks. **B.** PCA of normalized H3K27ac data, performed on all stitched enhancers (n of peaks = 17816). **C.** Heatmaps of normalized ATAC-seq peaks (left) and corresponding H3K27Ac signal (right) obtained from 3 normal beta cell donors and 4 insulinoma patients. Heatmaps are centered around ATAC-seq peak summits (±5 kb) and ranked by ATAC signal intensity. Peaks are classified into eight clusters (C1–C8) based on differential accessibility status and H3K27Ac enrichment status. C1 (n = 1,034), C3 (n = 613), and C6 (n = 6,808) represent beta-cell-enriched regulatory regions; C2 (n = 1,620) represents insulinoma-enriched regulatory regions; C4 (n = 1,922) and C5 (n = 19,180) represent shared regulatory regions. C7 (n = 15,323) and C8 (n = 62,410) were excluded from further analysis due to absence of H3K27Ac signal. C1, C3, and C6 were consolidated into Cluster 1; C2 into Cluster 2; and C4 and C5 into Cluster 3 in the main analysis (Fig. 1A).

**Supplementary Figure 2.**
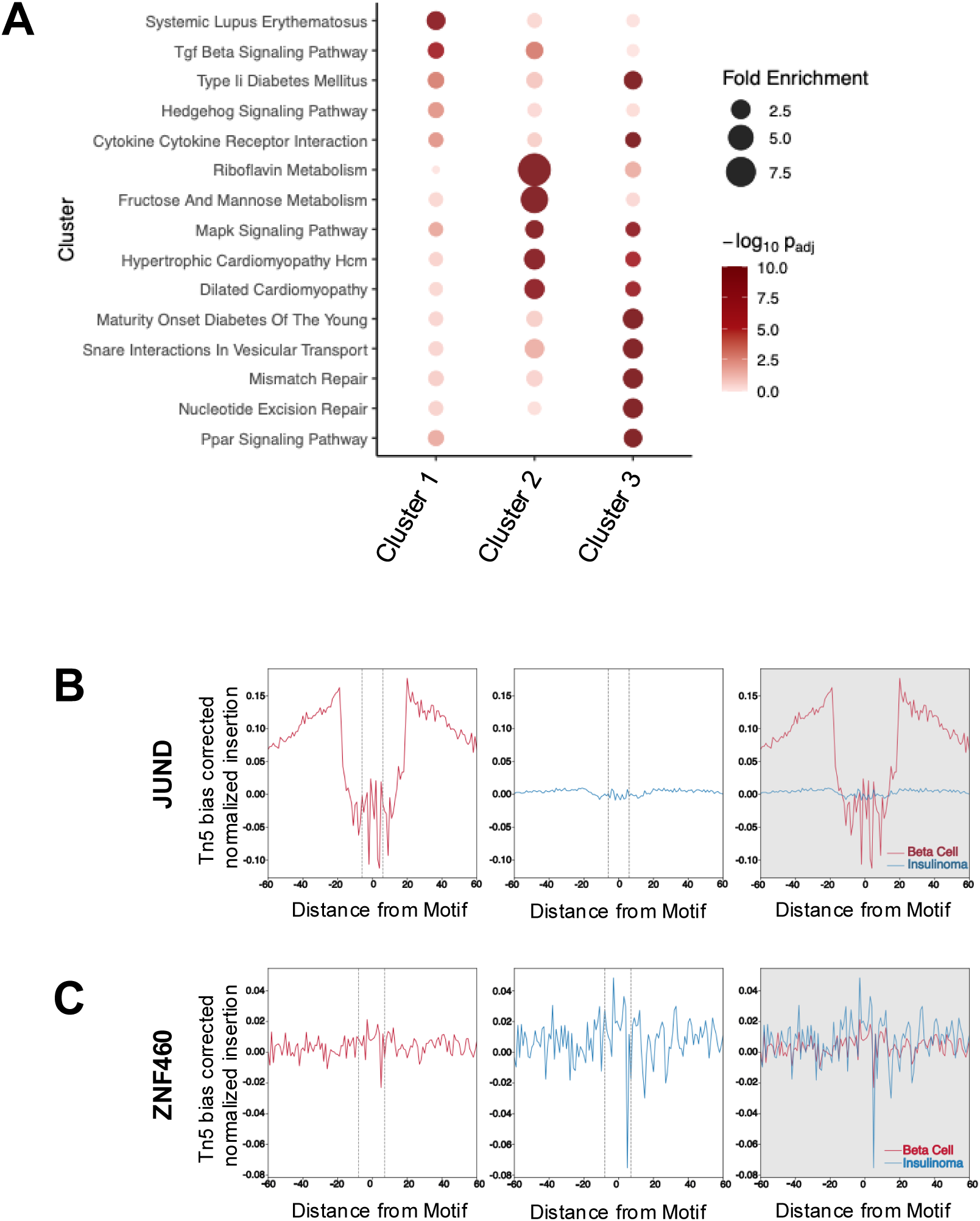
**A.** Enrichment of KEGG pathways for genes associated with beta cells, insulinomas or shared enhancers as in Fig 1A. The top 5 significant KEGG pathways for each cluster is shown. Dot size represents fold enrichment; color indicates statistical significance (−log10 adjusted P-value). Significance threshold: adjusted P<0.05. **B.** Transcription factor occupancy footprinting for JUND (n = 5,099 motif sites; top), ZNF460 (n = 1,882 motif sites; bottom). Plots show aggregated, Tn5 insertion-bias-corrected ATAC-seq signal (x-axis) ±60 bp centered on the motif center (y-axis). Signals from beta cell samples (n = 3) and insulinoma samples (n = 4) were merged within each group.

**Supplementary Figure 3.**
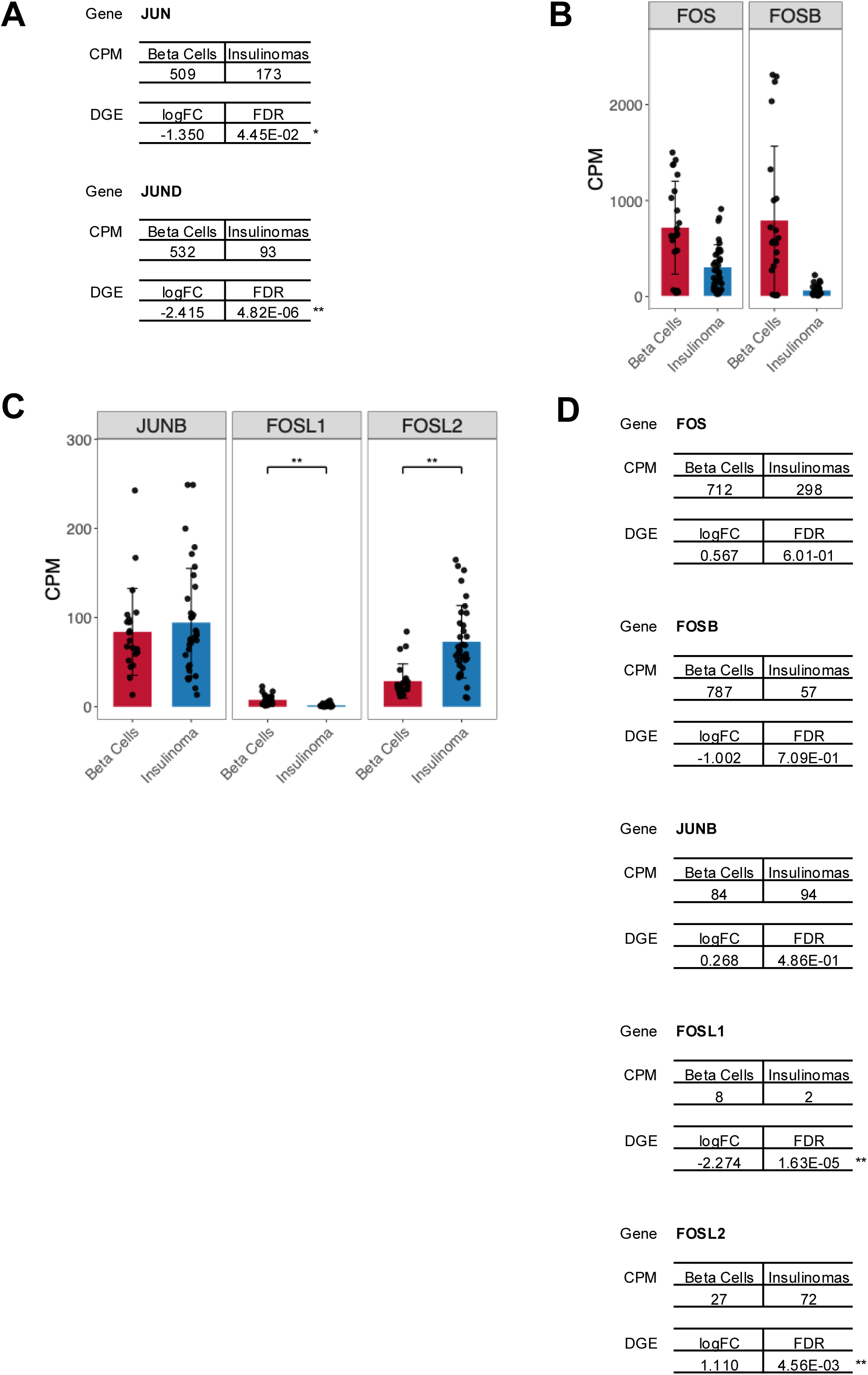
**A.** Summary of *JUN* and *JUND* bulk RNA-seq results, reporting mean CPM in beta cells and insulinomas and the corresponding differential expression statistics (logFC, FDR); significance is shown as *FDR < 0.05 and **FDR < 0.01. **B.** Bar chart of bulk RNA-seq expression (CPM) for AP-1 genes *FOS* and *FOSB* from differential gene expression analysis of 22 FACS-sorted beta cells (red) versus 34 insulinomas (blue). **C.** Bar chart of bulk RNA-seq expression (CPM) for AP-1 genes *JUNB*, *FOSL1* and *FOSL2* from differential gene expression analysis of 22 FACS-sorted beta cells (red) versus 34 insulinomas (blue). Bars indicate the group mean (±SD); dots represent individual samples. Brackets denote the between-group comparison; significance is shown as *FDR < 0.05 and **FDR < 0.01. **D.** Summary of *FOS*, *FOSB, JUNB*, *FOSL1* and *FOSL2* bulk RNA-seq results, reporting mean CPM in beta cells and insulinomas and the corresponding differential expression statistics (logFC, FDR); stars indicate the same FDR thresholds as in panel A.

**Supplementary Figure 4.**
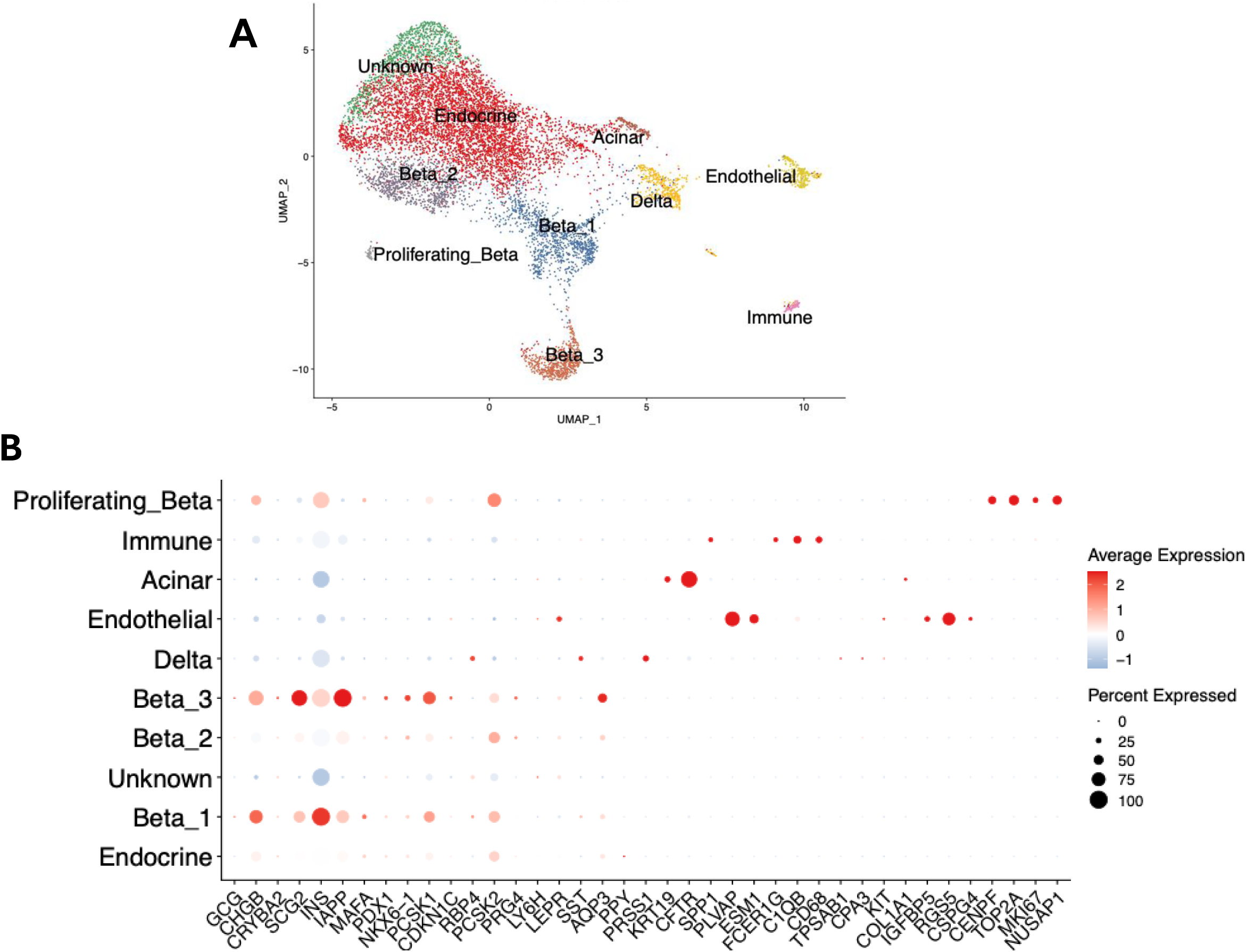
**A.** UMAP visualization of integrated snRNA-seq data from n = 3 insulinoma samples (9,730 cells in total), colored by inferred cell type. **B.** Dot plot showing the scaled normalized expression of canonical marker genes across cell type clusters used for annotation. Dot color represents the z-score of normalized mean expression; dot size indicates the percentage of cells within each cluster expressing the gene.

**Supplementary Figure 5.**
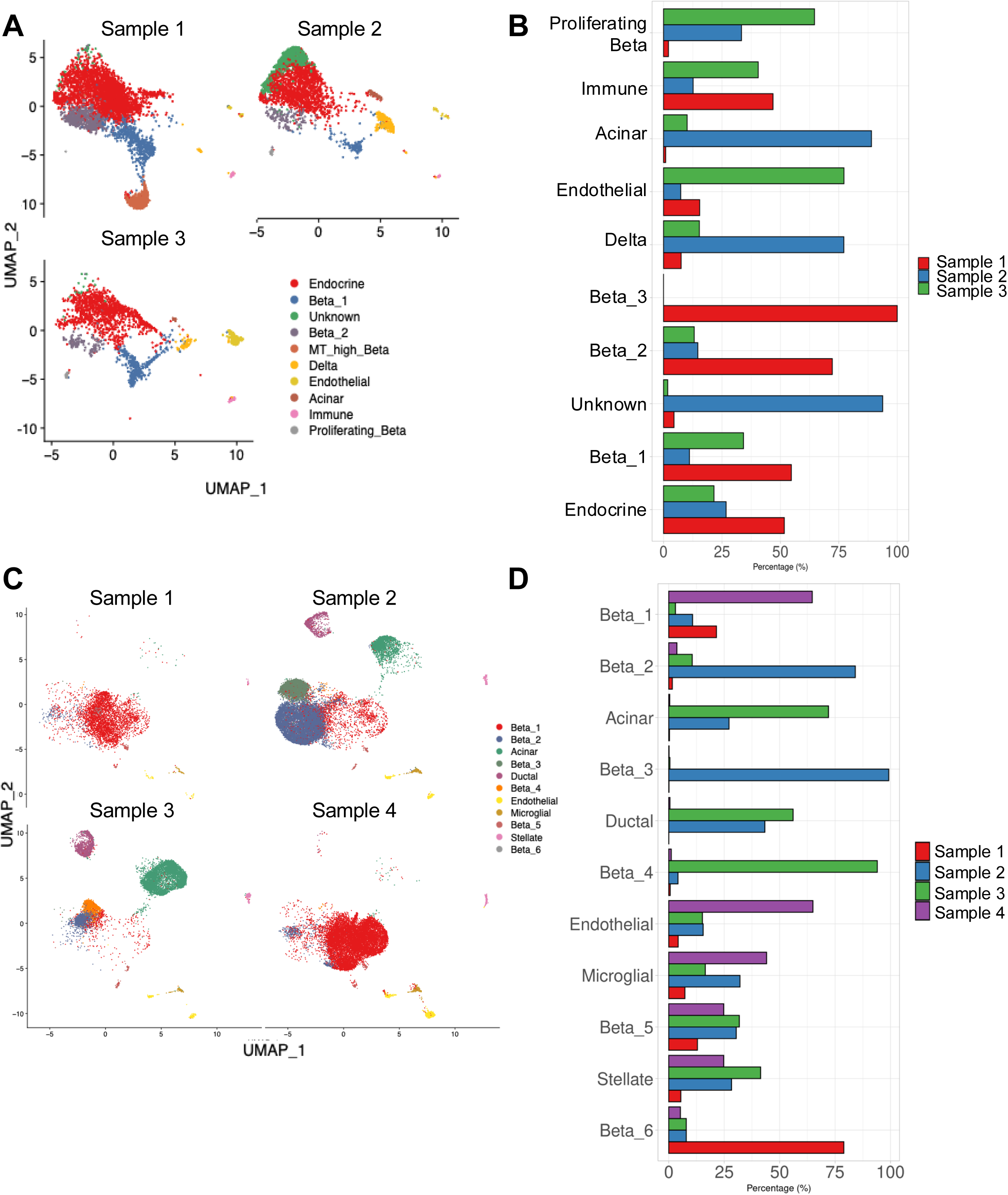
**A.** UMAP visualizations of integrated snRNA-seq data from n = 3 insulinoma samples displayed individually for each donor, colored by inferred cell type. Cell type labels follow the annotation described in Supplementary Fig. 4A. **B.** Stacked bar plot showing the percentage of cells from each individual insulinoma snRNA-seq sample (n = 3) assigned to each annotated cell type cluster. **C.** UMAP visualizations of integrated snATAC-seq data from n = 4 insulinoma samples displayed individually for each donor, colored by inferred cell type. Cell type labels follow the annotation described in Fig. 2A. **D.** Stacked bar plot showing the percentage of cells from each individual insulinoma snATAC-seq sample (n = 4) assigned to each annotated cell type cluster.

**Supplementary Figure 6.**
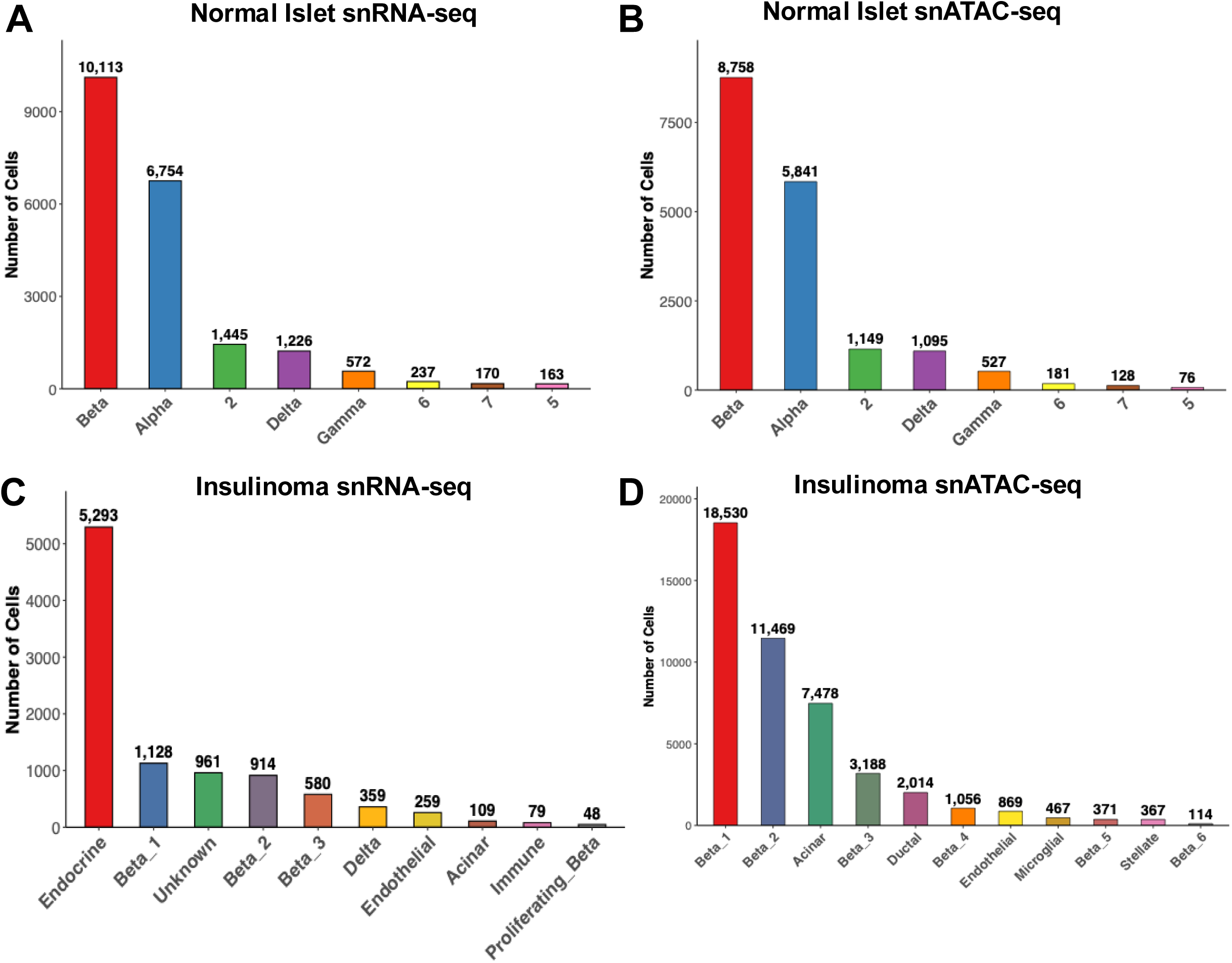
Bar plots showing the number of cells per assigned cell type in each dataset, sorted by abundance. **A.** Normal islet reference snRNA-seq (n = 20,680 cells). **B.** Normal islet reference snATAC-seq (n = 17,750 cells). **C.** Insulinoma snRNA-seq (n = 9,730 cells). **D.** Insulinoma snATAC-seq (n = 45,923 cells). Cell counts are displayed above each bar. Colors correspond to cell type assignments shown in the UMAP plots.

**Supplementary Figure 7.**
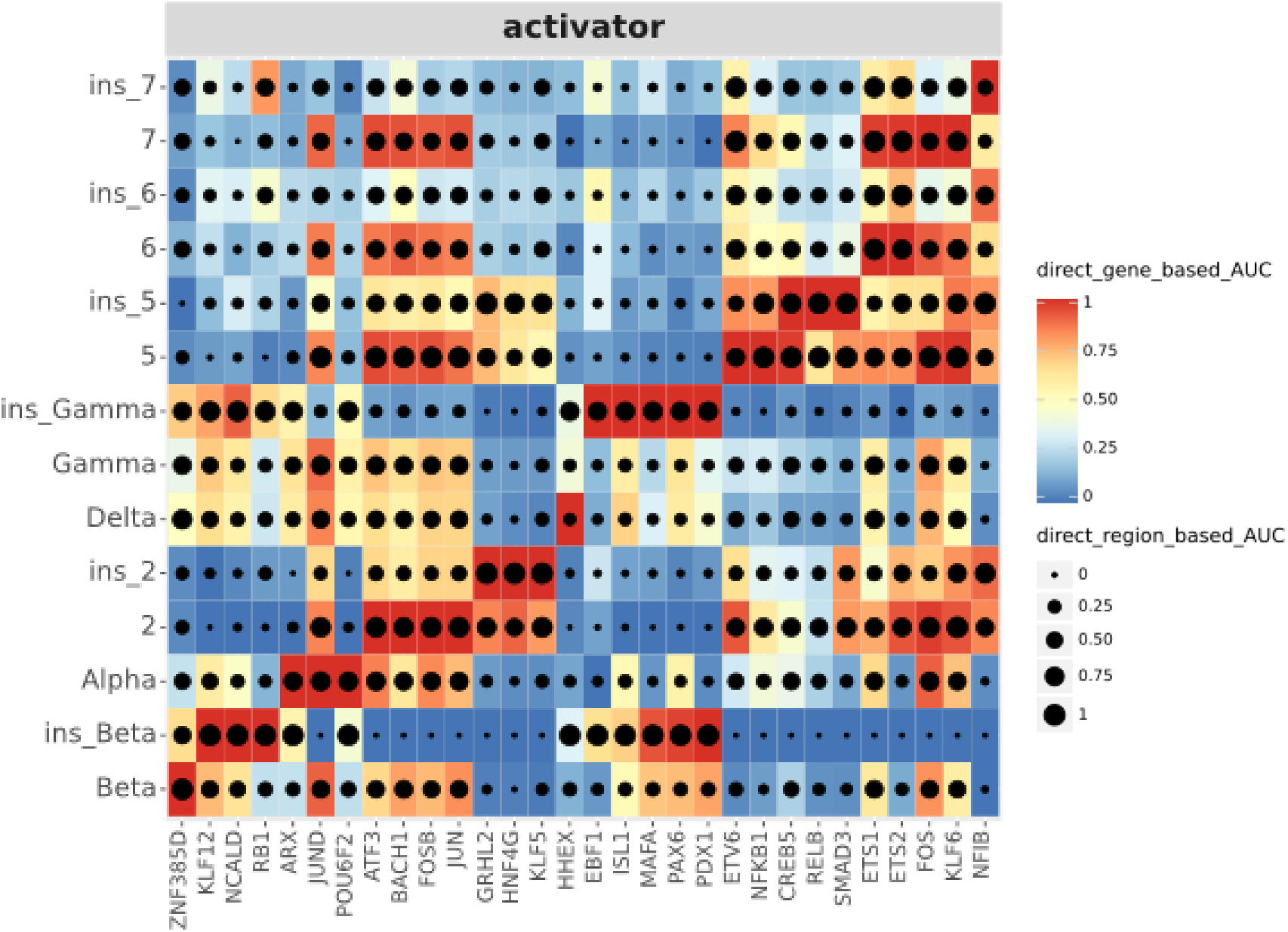
Heatmap/dot-plot showing the scaled AUC scores for the canonical activator transcription factor eRegulons within the top 50 most variable TF eRegulons. eRegulons are ordered by similarity of gene-based AUC profiles. Heatmap color indicates the expression-based AUC of target genes; dot size indicates the AUC of associated regulatory regions.

**Supplementary Figure 8.**
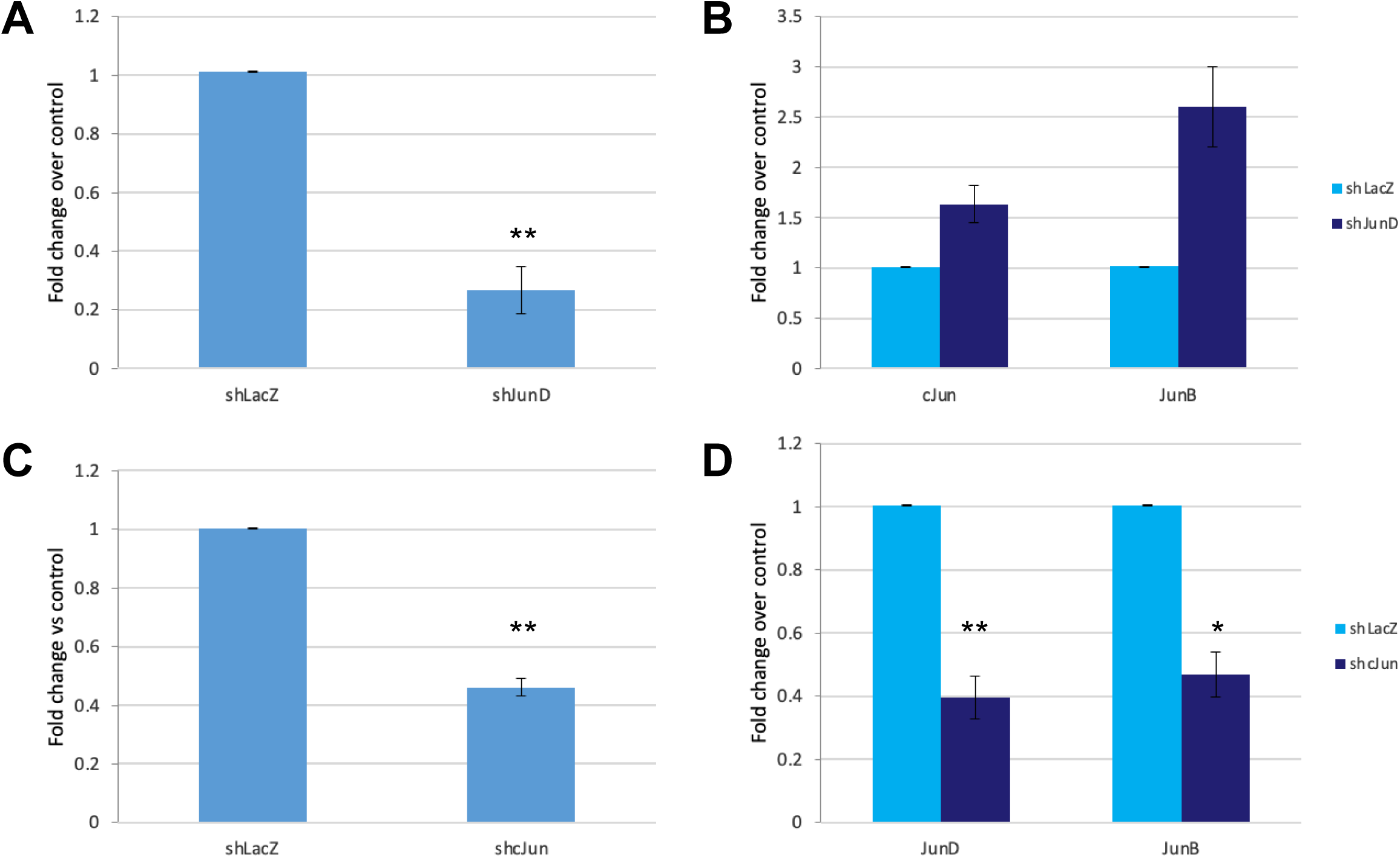
qRT-PCR results from dispersed human islets transfected with small hairpin RNAs targeting *JUN* or *JUND*. **A.** Gene expression level for *JUND* upon *JUND* silencing, confirming efficient silencing. **B.** Gene expression level for *JUN and JUNB* upon *JUND* silencing. Note that *JUN* and *JUNB* expression modestly increase with *JUND* silencing. **C.** Gene expression level for *JUN* upon *JUN* silencing, confirming efficient silencing. **D.** Gene expression level for *JUND* and *JUNB* upon *JUND* silencing. Note that *JUND* and *JUNB* expression levels are diminished when *JUN* is silenced.

**Supplementary Figure 9.**
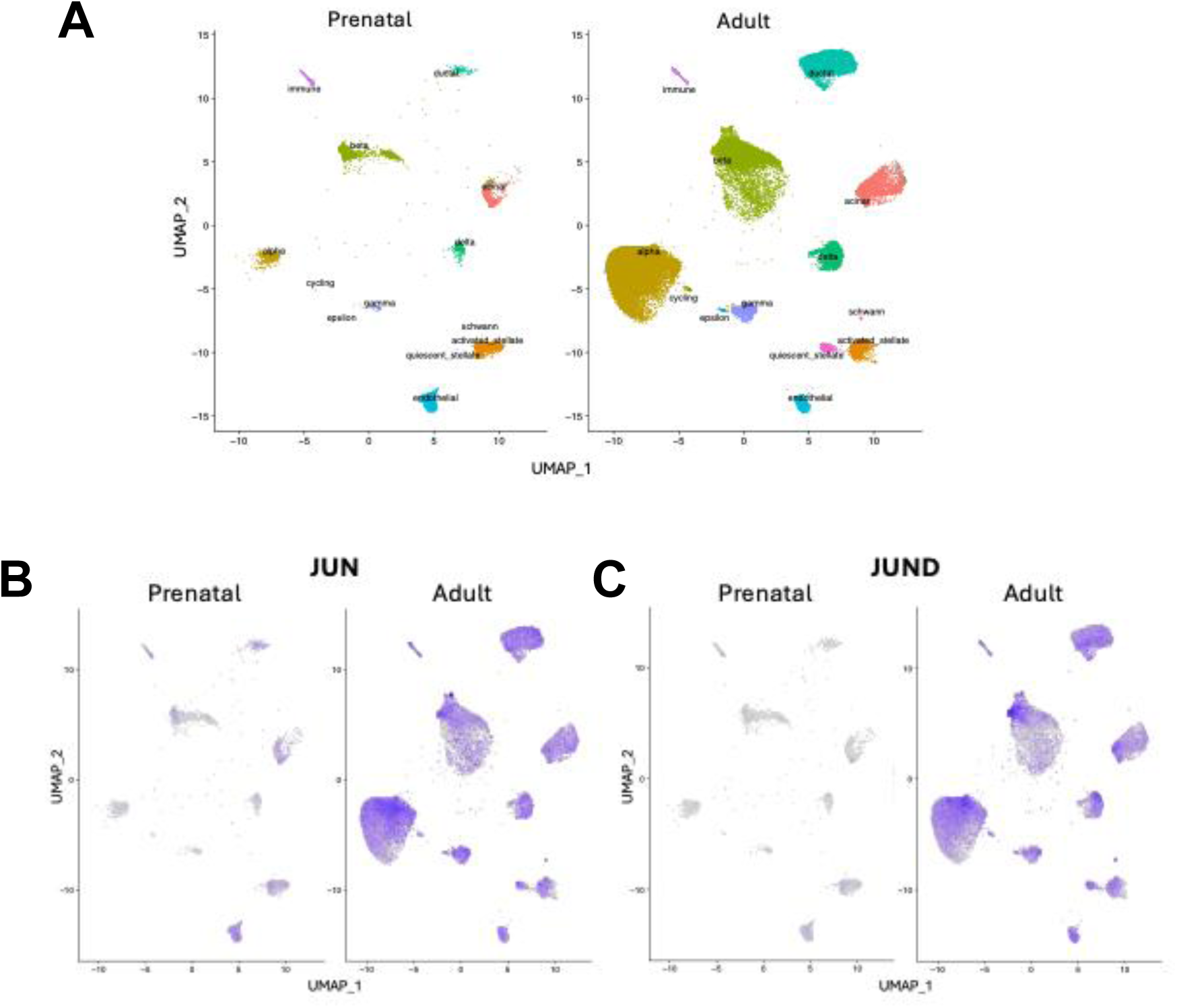
Integrated scRNA-seq atlas assembled from multiple publicly available datasets, showing prenatal and adult pancreatic cell populations after cross-study harmonization and reference-based annotation. **A.** Distribution of annotated cell types across the merged compendium, including endocrine, exocrine, stromal, endothelial, immune, and cycling populations. **B-C.** Feature plots for *JUN* and *JUND* across prenatal and adult cells. Signal is broadly detectable in both developmental stages, with strongest expression concentrated in adult endocrine and select non-endocrine compartments. Only cells passing annotation-confidence filtering were retained for feature plotting to improve interpretability of the multi-source integrated atlas.

**Supplementary Figure 10.**
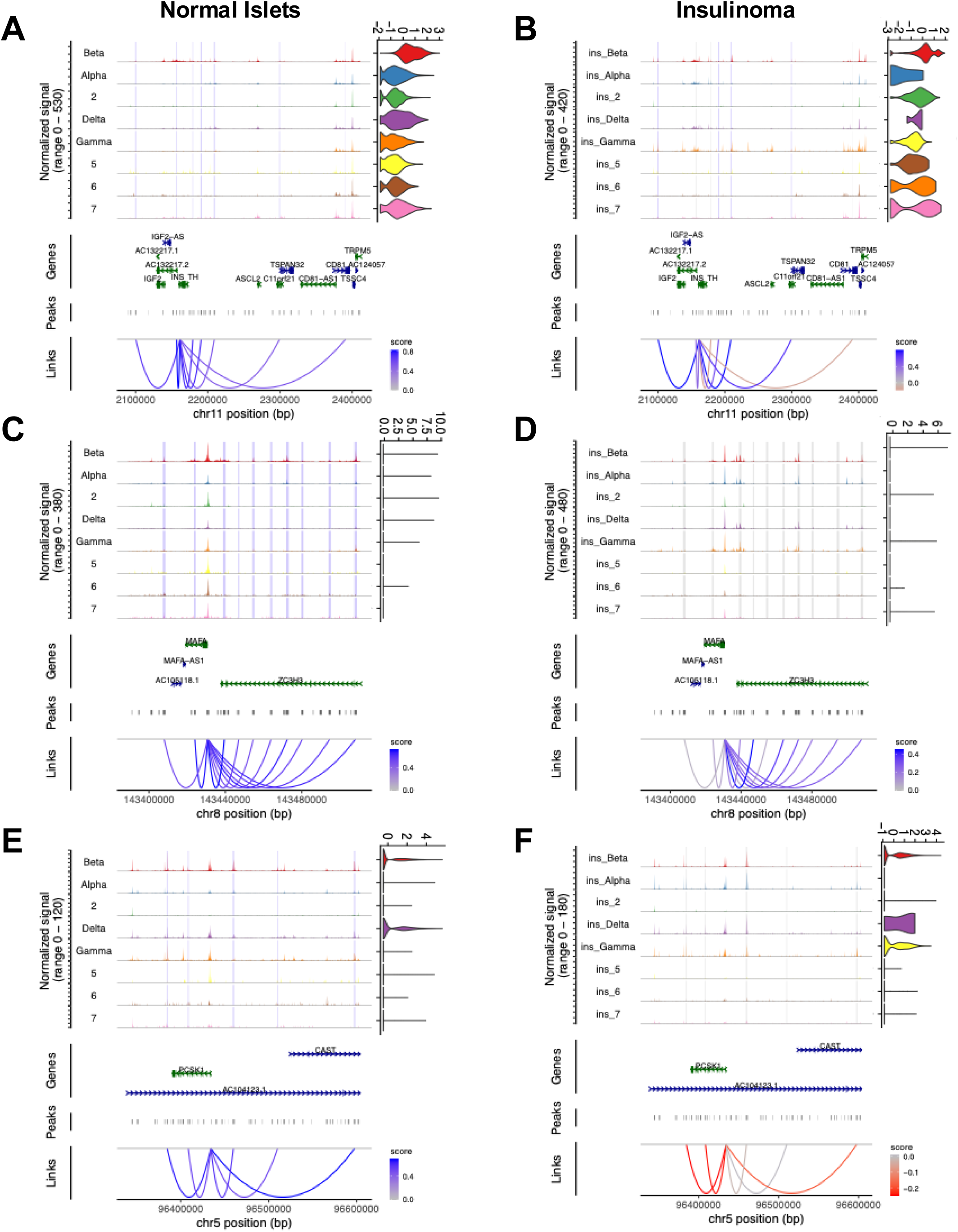
A-B. snATAC-seq coverage tracks at the *INS* locus for each cell cluster from normal islet (A) and insulinoma (B). ‘Normalized Signal’ track shows normalized, aggregated snATAC-seq signal from each cell type; minimum and maximum y-axis values are indicated adjacent to each track. Normalized expression of gene is shown as violin plot next to the genome browser track. Highlighted regions represent the distal regulatory elements found in cluster1 in **Fig5C** that significantly correlate with INS expression (blue) and insignificantly correlate with PDX1expresion (grey). Links from the significant distal regulatory elements to PDX1 TSS from cluster 1 (bottom). Note that these distal regulatory elements contain AP-1 binding motifs. Blue and red loops indicate positive and negative correlation with gene expression respectively. **C–D.** Shown for MAFA locus. **E–F.** Shown for PCSK1 locus.

