## Supplementary Table Information for "Epigenetic Profiling of Human Insulinomas Reveals AP-1 Family as Critical Regulators of Beta Cell Maturation"

**Supplementary Table 1** – ATAC-seq Peaks (supplementary_table_atac_peaks.xlsx)

Sheet1: Differential chromatin accessibility analysis of bulk ATAC-seq peaks between insulinoma and sorted beta cell samples. Differential test was performed using DESeq2.

Sheets2-4: ATAC peaks classified into three clusters through integrating with H3K27ac: beta cell-associated cluster1 (sheet2), insulinoma-associated cluster2 (sheet3), and shared cluster3 (sheet4).

Columns include genomic coordinates and statistics from differential accessibility test of each peak.

**Supplementary Table 2** - H3K27ac Enhancers (supplementary_table_h3k27ac_enhancers.xlsx)

Sheet1: Differentially enriched H3K27ac enhancer regions between insulinoma and sorted beta cell samples identified by DiffBind.

Sheet2-4: H3K27ac regions associated with cluster1 (sheet2), cluster2 (sheet3), and cluster3 (sheet4).

Columns include genomic coordinates and statistics from differential binding test of each peak.

**Supplementary Table 3** - rGREAT KEGG Pathway Enrichment (supplementary_table_rGREAT_KEGG.xlsx)

rGREAT pathway enrichment analysis using KEGG database for genomic regions in cluster1, cluster2, and cluster3. Genes that are detected in bulkRNAseq dataset are used as universe.

**Supplementary Table 4** - AME Motif Enrichment (supplementary_table_AME_motif_enrichment.xlsx)

Motif enrichment results in peaks associated with cluster1, cluster2 and cluster3 against ATAC-seq peaks that are not associated with H3K27Ac.

**Supplementary Table 5** - TOBIAS BINDetect (supplementary_table_tobias_bindetect.xlsx)

TOBIAS BINDetect transcription factor footprinting analysis within ATAC-seq peaks associated with cluster1 (sheet1), cluster2 (sheet2) and cluster3 (sheet3).

**Supplementary Table 6** - snRNA Cell Metadata (insulinoma_integrated_snrna_seurat_cell_metadta.csv)

Single-nucleus RNA-seq cell metadata for the insulinoma samples integrated through Seurat (n = 3).

**Supplementary Table 7** - snRNA Cluster Marker (supplementary_table_snrna_cluster_marker.csv)

Genes up-regulated in each cell cluster, identified by Wilcoxon rank-sum test comparing a given cluster against all other clusters.

**Supplementary Table 8** - snATAC Cell Metadata (insulinoma_integrated_cell_metadata.csv)

Single-nucleus ATAC-seq cell metadata for the integrated insulinoma dataset (n = 4). Harmony was applied to correct for batch effects.

**Supplementary Table 9**- Cluster-Specific Peaks (insulinoma_integrated_harmony_cluster_specific_peaks.csv)

Cluster-specific accessible chromatin peaks from the integrated single-nucleus ATAC-seq dataset.

**Supplementary Table 10**- chromVAR Motif Markers (insulinoma_integrated_harmony_chromvar_wilcox_cluster_markers.csv)

chromVAR transcription factor motif activity markers for each cell cluster, identified by Wilcoxon rank-sum test comparing a given cluster against all other clusters.

**Supplementary Table 11** - chromVAR Differential Motif Activity (chromvar_differential_activity_projected_id_broad_all_comparisons_merged.csv)

chromVAR differential motif activity results across pairwise cell type comparisons between insulinoma and normal islet in the insulinoma and normal islet integrated dataset. Harmony was applied to correct batch effect.

**Supplementary Table 12** - Peak-Gene Links with AP-1 Motifs (significant_links_with_ap1.csv)

Peak-to-gene linkage analysis with AP-1 motif annotation. Significant peak-gene links were identified separately in insulinoma and non-diabetic datasets by correlation of peak accessibility with gene expression.

**Supplementary Table 13** - HOMER De Novo Motif Enrichment (homer_de_novo_motif_results.xlsx)

HOMER de novo motif discovery results for cluster of peaks involved in the significant peak-to-gene links.

**Supplementary Table 14** - Demographic Information on Human Islet Donors

**Supplementary Table 15** - Demographic Information on Insulinoma Patients

**Supplementary Table 16** - List of Single Cell studies from Adult and Prenatal Donors

**Supplementary Table 17** - List of qRT-PCR Primer Sequences
